# Prrx1b directs pro-regenerative fibroblasts during zebrafish heart regeneration

**DOI:** 10.1101/2020.06.13.149013

**Authors:** Dennis E.M. de Bakker, Esther Dronkers, Mara Bouwman, Aryan Vink, Marie-José Goumans, Anke M. Smits, Jeroen Bakkers

## Abstract

**Rationale:** The human heart loses millions of cardiomyocytes after an ischemic injury, but is unable to regenerate the lost tissue. Instead, the injured human heart is repaired by pro-fibrotic fibroblasts that form a large permanent scar. In contrast, the injured zebrafish heart regenerates efficiently without the formation of a permanent scar. While fibroblasts have been shown to be indispensable for zebrafish heart regeneration, very little is known about the mechanisms balancing the fibrotic and regenerative response. A better understanding of these mechanisms could lead to the discovery of novel therapeutic strategies to reduce fibrosis and promote heart regeneration.

**Objective:** To identify novel mechanisms that regulate the balance between cardiac fibrosis and scar-free regeneration.

**Methods and Results:** Using a genetic approach, we first show that zebrafish *prrx1b* loss-of-function mutants display reduced cardiomyocyte proliferation and impaired heart regeneration. Using a lineage tracing approach, we show that Prrx1b is expressed in *tcf21*+ epicardial-derived cells localizing around and inside the injured area. Next, we used a single cell RNA-sequencing approach on sorted *tcf21*+ cells isolated from injured *prrx1b*-/- and wild-type hearts and identified two distinct fibroblast populations. With combined bioinformatic and histological analysis we found that *prrx1b*-/- hearts contain an excess of pro-fibrotic fibroblasts that produce TGF-β ligands and collagens, while fewer pro-regenerative Nrg1-expressing fibroblasts are formed. Furthermore, by injecting recombinant NRG1 in *prrx1b*-/- fish we were able to rescue their cardiomyocyte proliferation defect. Finally, using cultured human fetal epicardial cells and siRNA mediated knock-down of PRRX1 we found that PRRX1 is required for NRG1 induction in human epicardial-derived cells.

**Conclusions:** Prrx1b in the injured heart restricts fibrosis and stimulates regeneration by directing epicardial-derived cells towards a pro-regenerative Nrg1-producing fibroblast state.

## Introduction

Cardiovascular disease, including consequences of ischemic heart injury, still forms the leading cause of death in the western world (WHO 2019). Although most patients survive the initial injury, they lack an endogenous mechanism to regenerate the lost myocardium. Instead, a repair mechanism is activated in which the necrotic cardiomyocytes are removed and replaced by a fibrotic scar. While the fibrotic scar will preserve heart structure and functional integrity, it also induces pathological changes resulting in chamber dilatation, cardiomyocyte hypertrophy, apoptosis and ultimately heart failure (Baudino et al. 2006; Travers et al. 2016).

The fibrotic scar is formed by cardiac fibroblasts that become activated to produce large amounts of extracellular matrix (ECM) components, like collagens. Cardiac fibroblasts mainly originate from the embryonic epicardium, which consists of a heterogeneous population of cells (Acharya et al. 2012; Cao et al. 2016; Gittenberger-de Groot et al. 1998; Travers et al. 2016; Weinberger et al. 2020). During embryonic heart development a subset of epicardial cells undergoes an epithelial-to-mesenchymal transition (EMT) and migrate into the cardiac wall to give rise to a variety of cell types, which are commonly referred to as epicardial derived cells (EPDCs) and include predominantly cardiac fibroblasts and vascular support cells (Cao and Poss 2018).

In contrast to mammals, adult zebrafish can fully regenerate their heart after resection or cryoinjury of 20-30% of the ventricle (Chablais et al. 2011; González-Rosa et al. 2011; Poss, Wilson, and Keating 2002; Schnabel et al. 2011), which is due to the reactivation of proliferation in border zone cardiomyocytes (Jopling et al. 2010; Kikuchi et al. 2010; Wu et al. 2015). One of the first responses upon injury is the activation of a developmental gene expression program in the epicardium, at 1-2 days after the injury (Lepilina et al. 2006). This response becomes confined to the injury area as the epicardium is regenerated, which completely surrounds the injury area between 3 and 7 days post injury (dpi) (Kikuchi et al. 2011; Lepilina et al. 2006). The epicardium is essential for the regeneration process as ablating *tcf21*-expressing epicardial cells from the injured zebrafish heart impairs cardiomyocyte proliferation and regeneration (Wang et al. 2015). Similar to the results in the mammalian heart, lineage tracing of *wt1* and *tcf21* expressing cells in zebrafish revealed that the embryonic epicardium gives rise to EPDCs such as cardiac fibroblasts and vascular support cells (González-Rosa, Peralta, and Mercader 2012; Kikuchi et al. 2011; Sánchez-iranzo et al. 2018). Upon injury, EPDCs secrete signals guiding regeneration such as TGF-β ligands, platelet derived growth factor, cytokines like Cxcl12 and mitogenic factors such as Nrg1 (Chablais and Jazwinska 2012; Gemberling et al. 2015; Itou et al. 2012; Kim et al. 2010). In addition, EPDCs also contribute to fibrosis by producing ECM components such as collagen (Sánchez-iranzo et al. 2018). Fibrosis in the zebrafish heart is transient and regresses as regeneration proceeds, which coincides with the inactivation of the cardiac fibroblasts, ultimately resulting in a scar-free heart (Chablais et al. 2011; Sánchez-iranzo et al. 2018). While EPDCs play important roles during fibrosis and pro-regenerative signaling, the molecular mechanism regulating these processes remains largely unknown.

The paired related homeobox 1 (Prrx1) gene encodes a transcription factor and its expression correlates with scar-free wound healing and limb regeneration (Satoh et al. 2010; Stelnicki et al. 1998; Yokoyama et al. 2011). While its function was studied during embryonic development, its role during wound healing or regeneration remains largely unknown. Here we find that Prrx1b expression is induced in EPDCs during zebrafish heart regeneration and we show that Prrx1b is required for scar-free regeneration. By single cell RNA-sequencing of EPDCs we identify two distinct fibroblast subpopulations with either a pro-fibrotic or a pro-regenerative signature. Loss of Prrx1b results in an excess of pro-fibrotic fibroblasts and fibrosis, while fewer pro-regenerative fibroblast are formed. Furthermore, we find that Prrx1b stimulates cardiomyocyte proliferation by inducing Nrg1 expression in pro-regenerative fibroblasts and that this function is conserved in human fetal EPDCs. Altogether our data indicate that Prrx1b regulates the balance between pro-fibrotic and pro-regenerative fibroblasts to restrict fibrosis and stimulate cardiomyocyte proliferation during heart regeneration.

## Results

### *prrx1b* is required for zebrafish cardiomyocyte proliferation and heart regeneration

In order to address a potential role for Prrx1 in heart regeneration we utilized *prrx1a-/-* and *prrx1b-/-* adult fish, which harbour non-sense mutations upstream of the conserved DNA-binding domain (Barske et al. 2016). Both *prrx1a-/-* and *prrx1b-/-* display no developmental defects due to redundant gene functions and only *prrx1a-/-;/prrx1b-/-* embryos display craniofacial defects (Barske et al. 2016). We subjected adult *prrx1a-/-* and *prrx1b-/-* fish to cardiac cryoinjury and analysed scar sizes at 30dpi. While we observed comparable scar sizes in wild-type and *prrx1a-/-* hearts, scar sizes were significantly larger in *prrx1b-/-* hearts compared to their control siblings (**Fig.1A,B**, **Fig.S1A**). This difference in scar size was still apparent at 90dpi, when wild-type hearts had completely resolved their scars (**Fig.1C**,**D**). Since myocardial regeneration is achieved through the proliferation of the surviving cardiomyocytes at the injury border zone, we investigated cardiomyocyte proliferation in the *prrx1a-/-* and *prrx1b-/-* hearts. Indeed, *prrx1b*-/- hearts showed a significant reduction in border zone cardiomyocyte proliferation at 7dpi while no significant differences were observed in *prrx1a*-/- hearts **(Fig.1E,F, Fig.S2B,C)**. From these results we conclude that *prrx1b*, but not *prrx1a*, is required for zebrafish cardiomyocyte proliferation and heart regeneration.

### Prrx1 is expressed in epicardial and epicardial-derived cells

Next, to address how Prrx1 could function during zebrafish heart regeneration we investigated the spatial distribution of Prrx1 protein using an antibody raised against the N-terminus of the axolotl Prrx1 protein (Gerber et al. 2018). While Prrx1 was nearly undetectable in uninjured hearts, Prrx1 was clearly present in injured wild-type hearts and nearly absent from injured *prrx1b*-/- hearts (**Fig.S2A-C**). The localization of Prrx1 in the injury area suggested a possible expression in EPDCs. To validate this we used the *Tg(tcf21:CreERT2)* line which marks all EPDCs when crossed with the ubiquitous reporter *Tg(ubi:loxP-EGFP-loxP-mCherry)* and recombined during embryonic heart development (Kikuchi et al. 2011). Hearts from embryonic recombined *Tg(tcf21:CreERT2; ubi:loxP-EGFP-loxP-mCherry)* fish were cryo-injured, extracted at different time points and analysed for Prrx1 and mCherry expression (**Fig.2A-F**, **Fig.S3A-F**). At 1dpi, we observed a strong expression of Prrx1 in EPDCs around the entire ventricle (**Fig.2B**), coinciding with the previously reported ventricle-wide injury response of the epicardium (Lepilina et al. 2006). Importantly, while Prrx1 was mostly found in mCherry+ cells, not all mCherry+ cells were also positive for Prrx1, confirming the previously reported heterogeneity of EPDCs (González-Rosa et al. 2012). At 3dpi and 7 dpi, we started to observe double labelled mCherry+ and Prrx1+ cells around and in the injury area while Prrx1 expression became less apparent in the remote ventricle (**Fig.2C**,**D**). This resulted in very restrictive Prrx1 expression in a confined region in and around the injury area at 14dpi (**Fig.2F**). Similar to what we observed at earlier stages, Prrx1 expression was only detected in a subset of mCherry+ cells. At 30dpi, only a few mCherry/Prrx1+ cells were detected, of which the majority was located inside the remaining injury area. Taken together, these results indicate that upon cardiac injury, Prrx1 expression is induced in a subpopulation of EPDCs, which are initially localised at the epicardium and found more dispersed in the injury area at later stages.

### Proliferation and invasion of EPDCs is unaffected in *prrx1b-/-* hearts

Since cardiac injury induces the proliferation of EPDCs (Lepilina et al. 2006) and Prrx1b is expressed in this cell type we decided to investigate whether *prrx1b* plays a role in this process. To do so, we used the *Tg(tcf21:CreERT2; ubi:loxP-EGFP-loxP-mCherry)* line to identify EPDCs in wild-type and *prrx1b*-/- hearts and used PCNA expression to identify proliferating cells (**Fig.S4A**,**B**). We observed that the number of PCNA expressing EPDCs was high at 1 and 3 dpi, after which their number dropped to lower amounts from 7dpi onwards, which is in lines with previous reports (Lepilina et al. 2006). We observed no significant differences between wild-type and *prrx1b*-/- hearts in the number of PCNA expressing EPDCs at any of the examined time points (**Fig.S4B**).

Furthermore, we wondered whether *prrx1b* could play a role during the invasion of EPDCs into the injury area. To quantify this, we first measured the mCherry+ surface area in and around the injury area at different time points (**Fig.S4C**). Next, we determined which percentage of this total surface area is comprised of cells that have invaded into the injury area (**Fig.S4D**). We used this percentage as a measure of invasion efficiency of the EPDCs. At all timepoints, except for 3dpi, there was no significant difference in either the total measured area or percentage of mCherry+ cells inside the injury between the wild-type and *prrx1b-/-* hearts. At 3dpi, the percentage of mCherry+ cells invading the injury area is lower in the *prrx1b-/-* hearts, but this is completely mitigated from 7dpi onwards.

From these results we concluded that Prrx1 seems dispensable for proliferation of epicardial cells and EPDCs after injury. In addition, we found that invasion of EPDCs into the injury area does occur in *prrx1b-/-* hearts, albeit with a delay.

### Characterization of EPDCs in the regenerating heart by single cell sequencing

Next, we wanted to identify which processes are regulated by Prrx1b in EPDCs that could explain the impaired regenerative response observed in the *prrx1b*-/- hearts. Because the epicardium and EPDCs are formed by heterogenous cell populations, we decided to take a single-cell RNA-sequencing (scRNAseq) approach to characterize these cells within the context of heart regeneration. First, we isolated ventricles from recombined *Tg(tcf21:CreERT2; ubi:loxP-EGFP-loxP-mCherry)* wild-type sibling or *prrx1b-/-* fish at 7dpi and FACsorted the mCherry+ cells (**Fig.S5A-C)**. Next, we performed scRNAseq using the SORT-seq (SOrting and Robot-assisted Transcriptome SEQuencing) platform on the isolated single cells (Muraro et al. 2016) (**Fig.3A**,**B**). We analysed over 1,400 cells with >1000 reads per cell using the Race-ID3 algorithm, which resulted in the identification of 10 cell clusters grouped based on their transcriptomic features (**Fig.3C**,**D**, *Online supplementary Table 1*).

To confirm that the sorting strategy worked we plotted the expression of epicardial markers *tcf21, tbx18, aldh1a2* and *wt1b* and observed that most cells express at least one of these albeit in different patterns (**Fig.3E**). These differences in *tcf21, tbx18, aldh1a2* and *wt1b* expression confirm the previously observed heterogeneity of epicardial cells and EPDCs (Cao et al. 2015; Weinberger et al. 2020). To identify different cell types, we selected marker genes for known cell types and analysed their expression in the various clusters. The cells from clusters 4,7 and 10 express *tcf21* and *tbx18* but lack expression of *aldh1a2* and *wt1b*, suggesting that these represent epicardial cells covering the remote myocardium. Indeed, *in situ* hybridization (ISH) for additional genes with high expression in these clusters marks epicardial cells covering the uninjured part of the heart (**Fig.3F**). Vice versa, *wt1b* and *aldh1a2* are expressed in most of the remaining clusters, with highest expression in cluster 1. ISH for additional genes marking cluster 1 revealed that their expression was mostly located in the epicardial region covering the injury area but not the remote myocardium (**Fig.3G**).

Since the epicardium gives rise to fibroblasts and pericytes, we plotted known marker genes for these cell types. Pericytes express genes such as the potassium channel *kcnj8* and the Notch receptor *notch1b* (Guichet et al. 2015; Vanlandewijck et al. 2018). The expression of these genes is most abundant in cluster 6 and gene ontology analysis reveals cluster 6 has an enrichment in genes linked to smooth muscle contraction and cell junction organization. These findings suggest that cells in cluster 6 represent pericytes (**Fig.3I**). Periostin (*postnb*) and fibronectin (*fn1a*) are expressed in activated fibroblasts and both genes showed highest expression in clusters 2,3 and 9 (**Fig.3H**). In addition, we observed that these clusters are enriched for various other genes reported to be upregulated in activated fibroblasts (e.g. *dkk3b, fstl1b, ptx3*) (*Online supplementary Table 2*), suggesting that these clusters are formed by activated cardiac fibroblasts (Sánchez-iranzo et al. 2018). Gene ontology analyses with genes enriched in clusters 2,3 and 9 indeed indicated processes such as extracellular matrix organization, dissolution of fibrin clot, collagen biosynthesis and modifying enzymes (*Online supplementary Table 3*).

From these results we conclude that the scRNAseq results represent different populations of epicardial and epicardial-derived cells from the regenerating heart. Based on our analysis we divided the cells in four general groups: remote epicardium (clusters 4,7 and 10), injury epicardium (cluster 1), epicardial-derived fibroblasts (clusters 2, 3 and 9) and epicardial-derived pericytes (cluster 6).

Next, we looked at the lineage relation between the different cell clusters that were identified by scRNAseq. For this we used the RaceID3 algorithm, which was designed to construct lineage trees from scRNAseq data (Grün et al. 2016). This revealed that cells from clusters 4,7, and 10, representing remote epicardial cells, are directly linked to cells from cluster 1, which represents epicardial cells that cover the injury area (**Fig.3J**). Indeed, this corroborates the transgenic lineage tracing data that showed that the newly formed epicardium arises from the pre-existing epicardium. Interestingly, the lineage tree suggests that cells from cluster 1 are linked to cluster 5, from which the lineage tree splits towards the pericyte cluster and the fibroblast clusters. This suggests that cells in cluster 5 represent a precursor cell type and that from here, cells differentiate into the two main cell types: pericytes (cluster 6) and fibroblasts (cluster 2,3). The pericytes and fibroblasts expressed *snai2* and *snai1*, two EMT inducers that are expressed in EPDCs when migrating into the cardiac wall (**Fig.3K**) (Barrallo-Gimeno and Nieto 2005). Together, the lineage analysis shows the interrelation between the various epicardial and epicardial derived clusters, suggesting that a subset of epicardial cells undergo EMT to form EPDCs that differentiate into fibroblasts and pericytes. Moreover, the cluster and lineage tree analysis suggest that there are two distinct fibroblasts clusters, cluster 2 and 3, that arise from cluster 5 separately.

### Prrx1 represses fibrosis and promotes *nrg1*-expressing EPDCs

Next, we mapped the wild-type and *prrx1b*-/- cells separately on the t-SNE map to reveal differences in contribution to the various cell clusters (**Fig.4A**,**B**). Interestingly, we found a clear difference in the contribution of wild-type and *prrx1b*-/- cells to the fibroblast clusters 2 and 3. While cluster 2 consists of mostly wild-type cells (89%) and few *prrx1b-/-* cells (11%), cluster 3 is enriched in *prrx1b-/-* cells (71%) compared to wild-type cells (29%) (**Fig.4B-D**). While both cluster 2 and cluster 3 cells express markers for activated fibroblasts such as *postnb*, the cluster and lineage analysis indicate transcriptional differences between cluster 2 and cluster 3 cells. Differential gene analysis between cluster 2 and 3 revealed that cluster 3 cells are enriched for genes involved in TGF-β signaling (*tgfb1a, tgfb2, tgfb3, P*.*val: 7*.*1E-4*), extracellular matrix proteins (*P*.*val: 3*.*3E-4*) and collagen fibril organization (*P*.*val: 1*.*2E-4*) (**Fig. 4E**). This suggests that cluster 3 cells are pro-fibrotic fibroblasts that produce collagen and other ECM proteins and that these are more abundant in *prrx1b-/-* hearts. To validate these findings, we performed ISH for genes with high expression in cluster 3 cells. Indeed, we observed increased expression of *tgfb1a* and *col11a1b* in injured *prrx1b*-/- hearts compared to their wild-type siblings (**Fig.4F**,**G**). In addition, we performed Sirius Red staining to visualize collagen, which showed an excess of collagen fibres in the *prrx1b*-/- hearts in and around the injury area (**Fig.4H-J**). Form these results we conclude that in injured *prrx1b-/-* hearts an excess of TGF-β ligand and ECM producing fibroblasts is formed, resulting in an enhanced fibrotic response to the injury.

**Figure 1:**
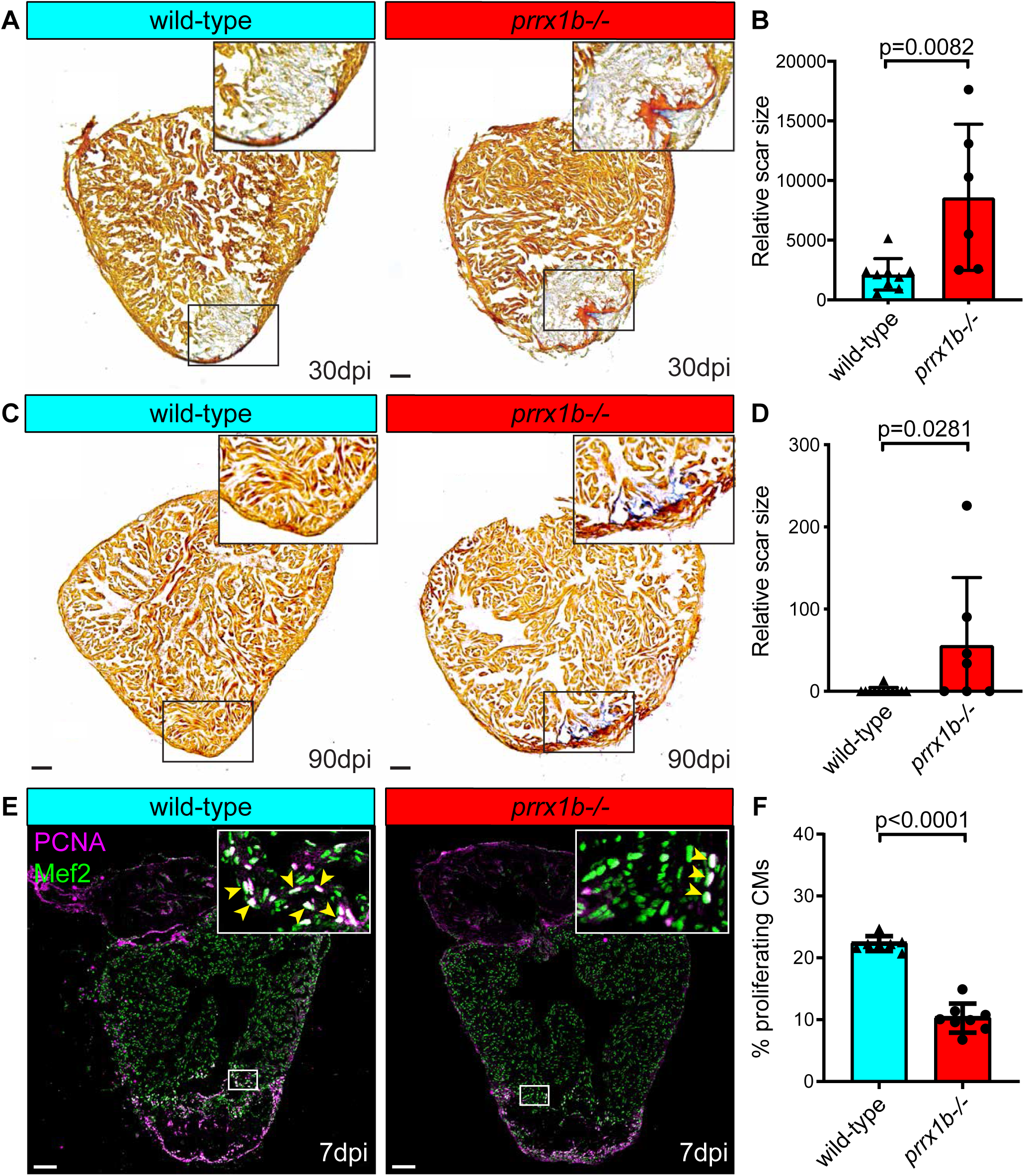
Heart regeneration and border zone cardiomyocyte proliferation is impaired in *prrx1b*-/- zebrafish. (**A**) AFOG staining on 30dpi wild-type and *prrx1b*-/- heart sections showing fibrin in red, collagen in light blue and remaining muscle tissue in orange. Scale bars represent 100μm. (**B**) Quantification of the remaining scar size at 30dpi between *prrx1b*-/- hearts and wild-type siblings. (**C**) AFOG staining on 90dpi wild-type and *prrx1b*-/- heart sections. Scale bars represent 100μm. (**D**) Quantification of the remaining scar size at 90dpi between *prrx1b*-/- hearts and wild-type siblings. Error bars represent SD. (**E**) Immunofluorescent staining on 7dpi wild-type and *prrx1b*-/- heart sections using an anti-Mef2 antibody as a marker for cardiomyocyte nuclei, and an anti-PCNA antibody as a nuclear proliferation marker. Arrowheads in zoom-ins indicate proliferating cardiomyocytes. Scale bars represent 100μm in the overview images and 10μm in the zoom-ins. (**F**) Quantification of the percentage of (PCNA+) proliferating border zone cardiomyocytes between *prrx1b*-/- hearts and their wild-type siblings. Error bars represent SD.

**Figure 2:**
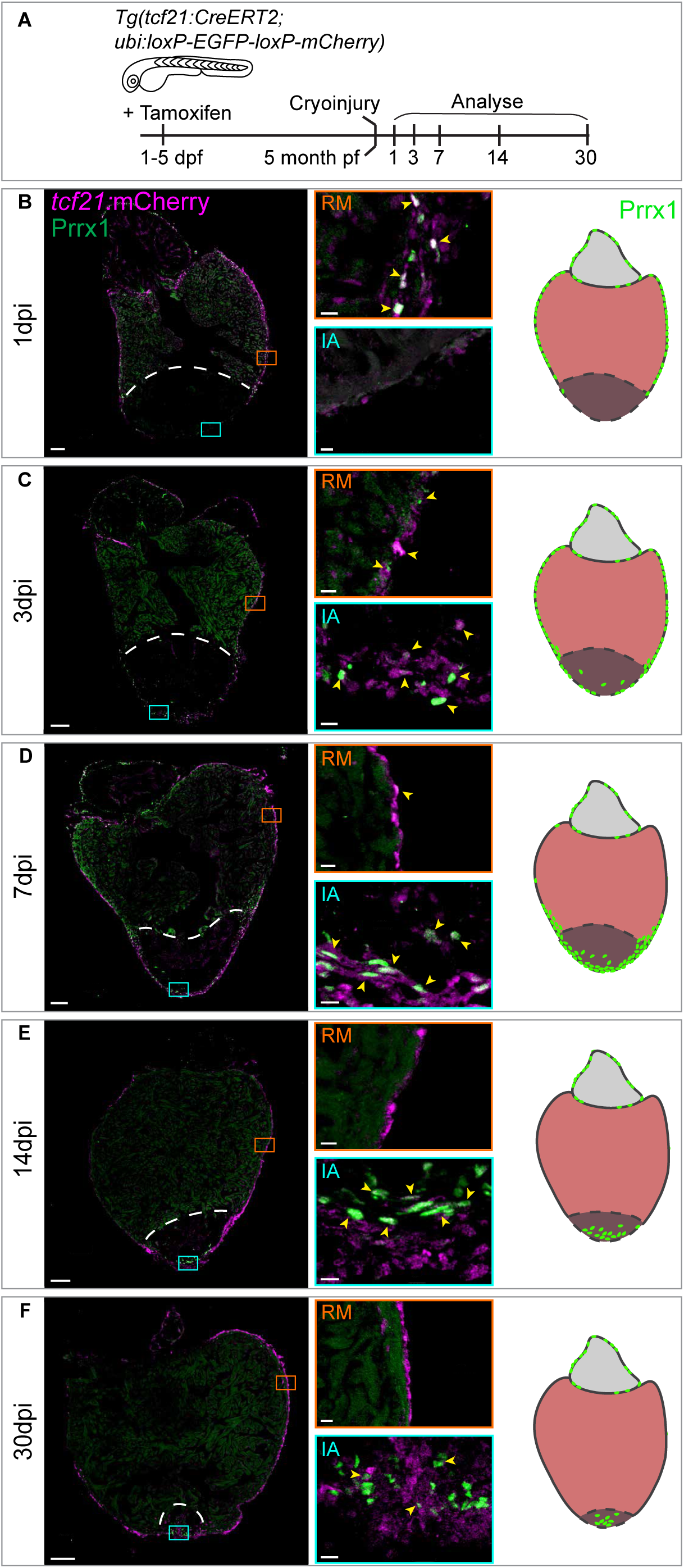
Prrx1 is expressed in EPDC and follows epicardial dynamics post injury. (**A**) Cartoon illustrating experimental procedures. (**B-F**) (Left panels) Immunofluorescent staining on uninjured, 1, 3, 7, 14 and 30dpi wild-type heart sections. Immunofluorescent staining on *tcf21*:mCherry+ wild-type heart sections using an antibodies against Prrx1 (green) and mCherry (magenta) to mark *tcf21*:mCherry+ cells. Dotted line represents injury border. Arrowheads indicate *tcf21*:mCherry+/Prrx1+ cells. Scale bars represent 100μm in the overview images and 10μm in the zoom-ins. RM= Remote myocardium, IA= Injury area. Hearts analysed per condition: 6. (Right panels) Schematic representation of Prrx1 dynamics upon injury. Prrx1+ cells are in green. Dark color at the apex represents the injury area.

**Figure 3:**
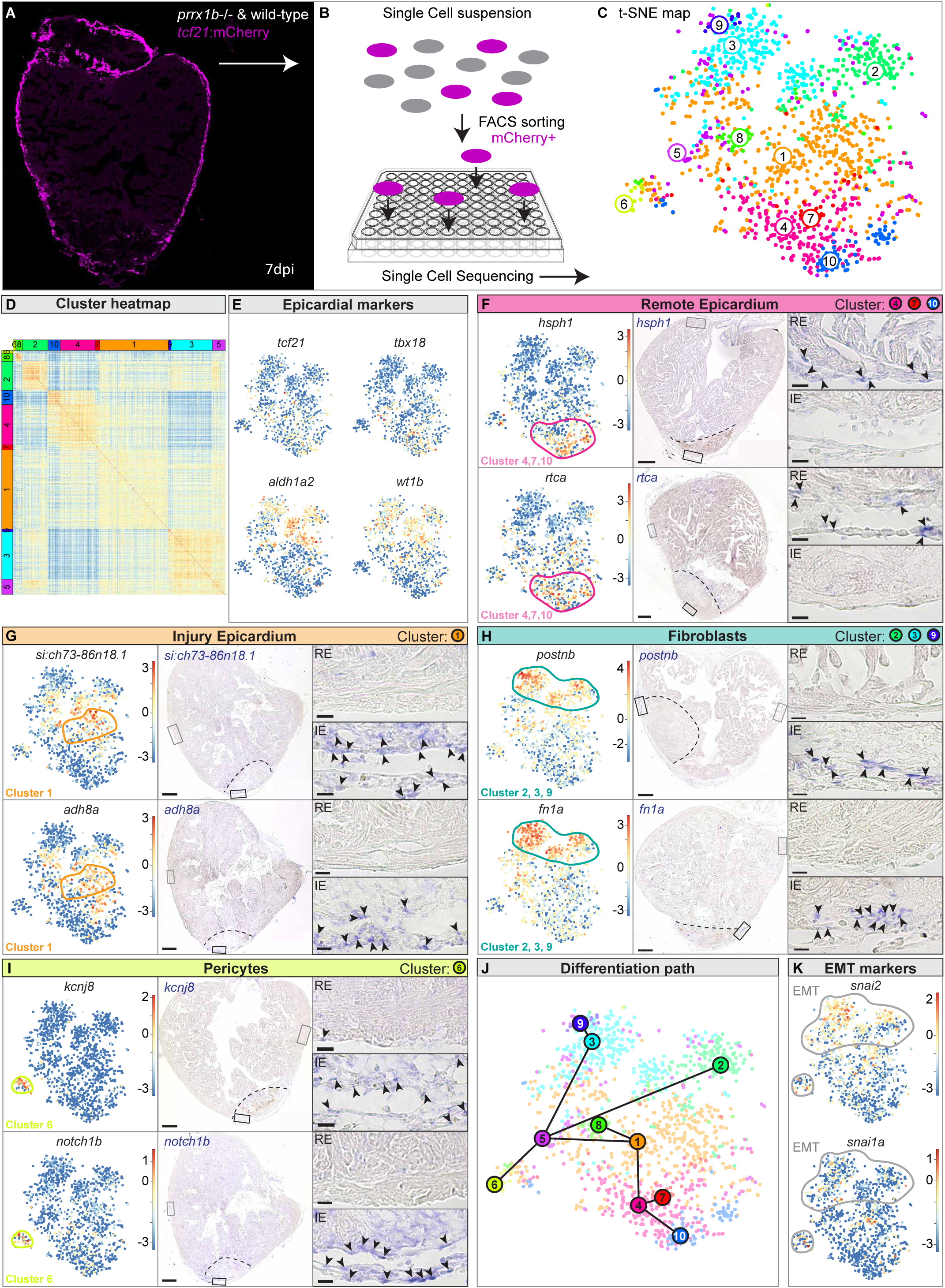
Single cell sequencing identifies epicardial derived cell populations in the injured zebrafish heart. (**A**) Workflow of the isolation and (**B**) sorting of tcf21:mCherry+ cells out of wildtype and *prrx1b*-/- hearts at 7dpi. (**C**) tSNE-plotting of the data results in 10 transcriptionally distinct clusters, as also indicated by the heatmap (**D**). (**E**) tSNE maps visualizing log2-transformed read-counts for *tcf21, tbx18, aldh1a2* and *wt1b*. (**F-I**) Characterization of the different cell clusters. Left panels show tSNE maps visualizing log2-transformed read-counts for genes with high expression in the indicated cluster. Middle panels display *in situ* hybridization for the cluster-enriched genes in wild-type hearts at 7 dpi. Dashed line indicates injury border and scale bars represent 100μm. Right panels display magnifications of the boxed regions in remote (RM) and injury area (IA) with arrowheads pointing to cells with high expression. Scale bars represent 10μm. Hearts analysed per condition: 3 (**J**) Inferred lineage tree of epicardial cells. tSNE map with the (significant) links between the cell clusters. (**K**) tSNE maps visualizing log2-transformed read-counts for *snai2* and *snai1a*. Gene lists are provided in *Online supplementary table 1*.

**Figure 4:**
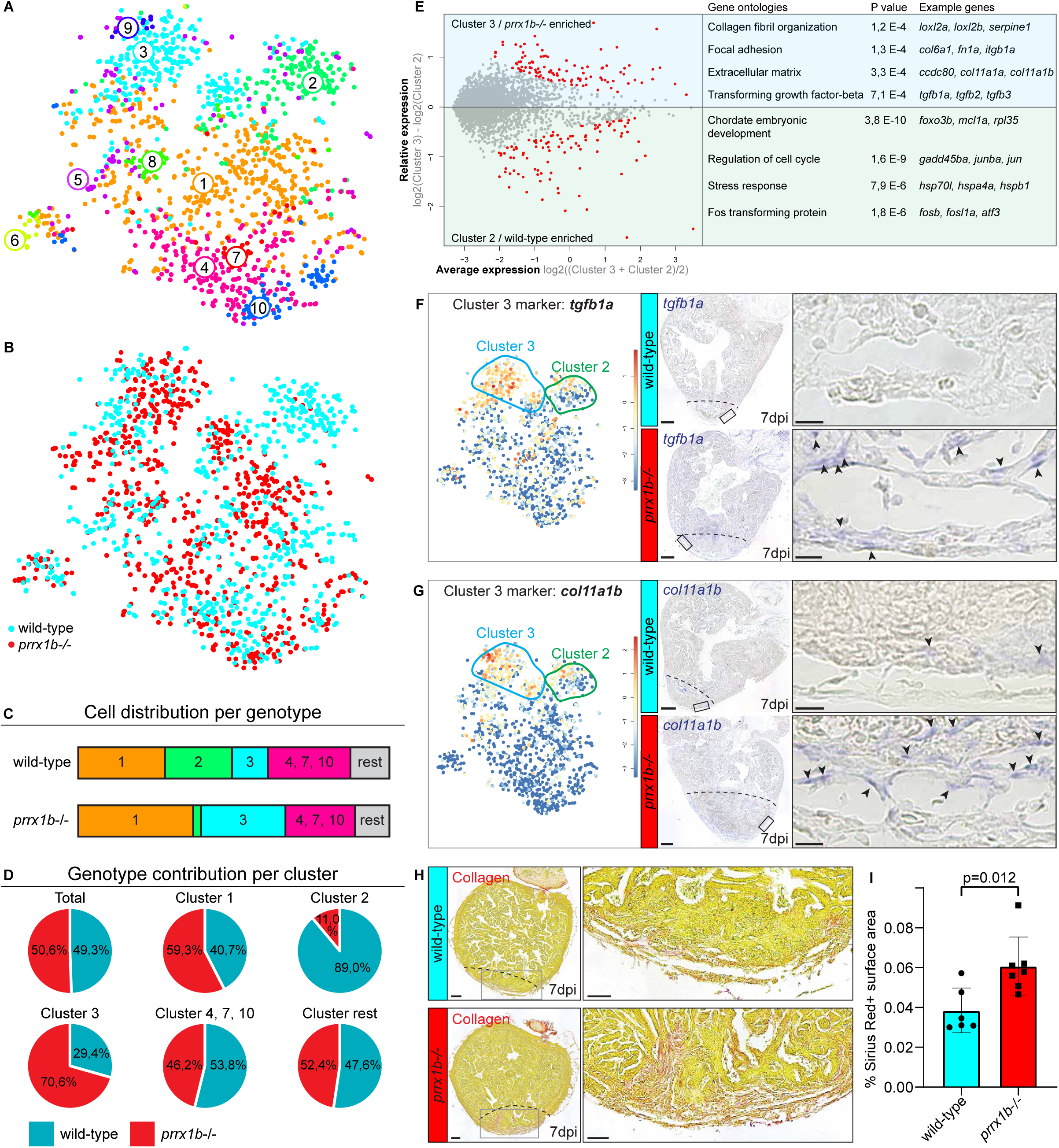
*prrx1b-/-* hearts contain excussive amounts of pro-fibrotic fibroblasts. (**A**) tSNE-map of the single cell sequencing data, indicating 10 transcriptionally distinct cell populations. (**B**) tSNE-map showing the contribution of wild-type cells (cyan) and *prrx1b*-/- cells (red). (**C**) Stacked bar graph showing relative cell contribution to major clusters in either wild-type or *prrx1b*-/- hearts. (**D**) Pie charts showing contribution of wild-type and *prrx1b*-/- cells per cluster. (**E**) Differential gene expression analysis using the DESeq algorithm between fibroblast clusters 2 and 3. Enriched genes were selected for either cluster 2 or 3 with a P-value cut-off of <0.05 (red). Gene ontology analysis was performed using the online tool DAVID. Gene and full Gene Ontology lists are provided in *Online supplementary table 2 and 3*. (**F-G**) Characterization of cluster 3. Left panels show tSNE maps visualizing log2-transformed read-counts for genes with high expression in the indicated cluster. Middle panels display *in situ* hybridization for the cluster 3-enriched genes in wild-type and *prrx1b*-/- hearts at 7 dpi. Dashed line indicates injury border and scale bars represent 100μm. Right panels display magnifications of the boxed regions in injury area with arrowheads pointing to cells with high expression. Scale bars represent 25μm. Hearts analysed per condition: 3 (**H**) Sirius red staining showing collagen in red on sections of wild-type and *prrx1b*-/- hearts at 7 dpi. Middle and right panels show magnifications of boxed regions in sub epicardial layer and further inside the injury area. Scale bars represent either 100μm (overview) or 50μm (zoom-in). (**I**) Quantification of Sirius Red (collagen) staining in wild-type and *prrx1b*-/- mutant hearts showing significantly more fibrosis in *prrx1b*-/- hearts inside and around the injury area.

While the observed excess of collagen production in injured *prrx1b-/-* fish could explain the larger scars at later time points, it is less likely an explanation for the reduced cardiomyocyte proliferation that was observed at 7dpi (**Fig.1E**,**F**). Both EPDCs and fibroblasts are known to secrete growth factors that induce cardiomyocyte proliferation (Gemberling et al. 2015; Ieda et al. 2009; Wang et al. 2015). Therefore, we hypothesized that cluster 2 cells, which are almost completely absent in *prrx1b-/-* hearts, produce a secreted growth factor that stimulates cardiomyocyte proliferation. Neuregulin (Nrg1) is a good candidate since it is expressed by EPDCs and can efficiently stimulate cardiomyocyte proliferation (D’Uva et al. 2015; Gemberling et al. 2015). We first addressed whether *nrg1* expression is enriched in cluster 2 cells, but *nrg1* expression is too low in the scRNAseq data to draw any conclusions (<100 combined reads from 1438 cells). We therefore investigated expression of *nrg1* through ISH and observed that in the epicardium expression of *nrg1* overlaps with the expression of *si:dkeyp-1h4*.*9*, a marker for cluster 2 cells (**Fig.S6**). Consistent with this finding, we observed that while the expression of both *nrg1* and *si:dkeyp-1h4*.*9* was prominent in the epicardial and sub-epicardial region in wild-type hearts at 7dpi, their expression was nearly undetectable in injured *prrx1b*-/- hearts (**Fig.5A**,**B**).

The observed reduction in *nrg1*-expressing fibroblasts in *prrx1b-/-* hearts could explain the reduction in proliferation of border zone cardiomyocyte. To test this hypothesis, we attempted to rescue the cardiomyocyte proliferation defect in *prrx1b-/-* hearts by daily intraperitoneal injection of recombinant NRG1 protein from 3dpi to 7dpi. Importantly, injecting *prrx1b*-/- zebrafish with recombinant NRG1 protein indeed restored cardiomyocyte proliferation in the border zone to wild-type levels (**Fig.5C**,**E**).

Together, these results indicate that Prrx1b restricts the number of pro-fibrotic epicardial-derived fibroblasts and stimulates the differentiation of *nrg1*-expressing, epicardial-derived fibroblasts that stimulate cardiomyocyte proliferation during zebrafish heart regeneration (**Fig.7**).

### PRRX1 regulates NRG-1 expression in human fetal epicardial cells

To place our findings in a broader context, we set out to investigate the relevance of PRRX1 to the human heart. Epicardial activation upon cardiac injury is conserved amongst vertebrate species. Following myocardial infarction, the murine epicardium quickly activates an embryonic gene program resulting in the re-expression of *Tbx18* and *Wt1* and *Tcf21* (Huang et al. 2012; Limana et al. 2010; van Wijk et al. 2012; Zhou et al. 2011). While in the mammalian neonatal heart this epicardial activation is followed by cardiomyocyte proliferation and regeneration, the adult mammalian heart shows only severely limited cardiomyocyte proliferation and forms a permanent scar instead (Porrello et al. 2011). Furthermore, fetal EPDCs play a crucial role in cardiac development which is party through the regulation of cardiomyocyte proliferation (Chen et al. 2002; Gittenberger-De Groot et al. 2000; Lavine et al. 2005; Li et al. 2011a). Therefore, we wondered whether PRRX1 could play a similar role in human fetal EPDCs as observed in the regenerating zebrafish heart. To obtain human fetal EPDCs we exploited a previously set-up *in vitro* model (Dronkers et al. 2018) in which human fetal epicardial cells can be cultured in an epithelial phenotype in the presence of the ALK4/5/7 kinase inhibitor SB431542. Removal of the inhibitor for at least 5 days results in the induction of EMT, which can be appreciated by the transition of cobblestone epithelial-like cells towards spindle-shaped mesenchymal cells and upregulation of mesenchymal genes *POSTN* and *FN1* (**Fig.6A-C**) (Moerkamp et al. 2016). Although some *PRRX1* expression was detected in cobblestone epithelial-like cells, its expression was significantly increased in spindle-shaped mesenchymal cells (**Fig.6C**). Interestingly, *NRG1* expression followed the same pattern as *PRRX1* expression, as they were both increased in spindle-shaped cells. To determine whether PRRX1 is required for *NRG1* expression as found during zebrafish heart regeneration, spindle-shaped mesenchymal cells were subjected to *PRRX1* knock-down (KD) using siRNAs (**Fig.6D**). Indeed, *PRRX1* KD led to a significant decrease in *NRG1* mRNA, as well as a significant decrease of secreted NRG1-β1 protein (**Fig.6E**,**F**).

Next, we aimed to investigate the relevance of PRRX1 during human heart repair and analysed PRRX1 expression in human tissue from eight patients that suffered a recent myocardial infarction, showing beginning scar formation and fibroblasts infiltration (**Fig. 6G**). Positive staining could be observed in human colon and lung tissue, which indicates that the staining protocol is able to detect PRRX1 in human tissue (**Fig.6H**,**I**). However, only very few PRRX1+ cells could be identified in human injured heart tissue, either in the (sub-)epicardium, sub-epicardial myocardium or in the infarcted area (**Fig.6J**).

**Figure 5:**
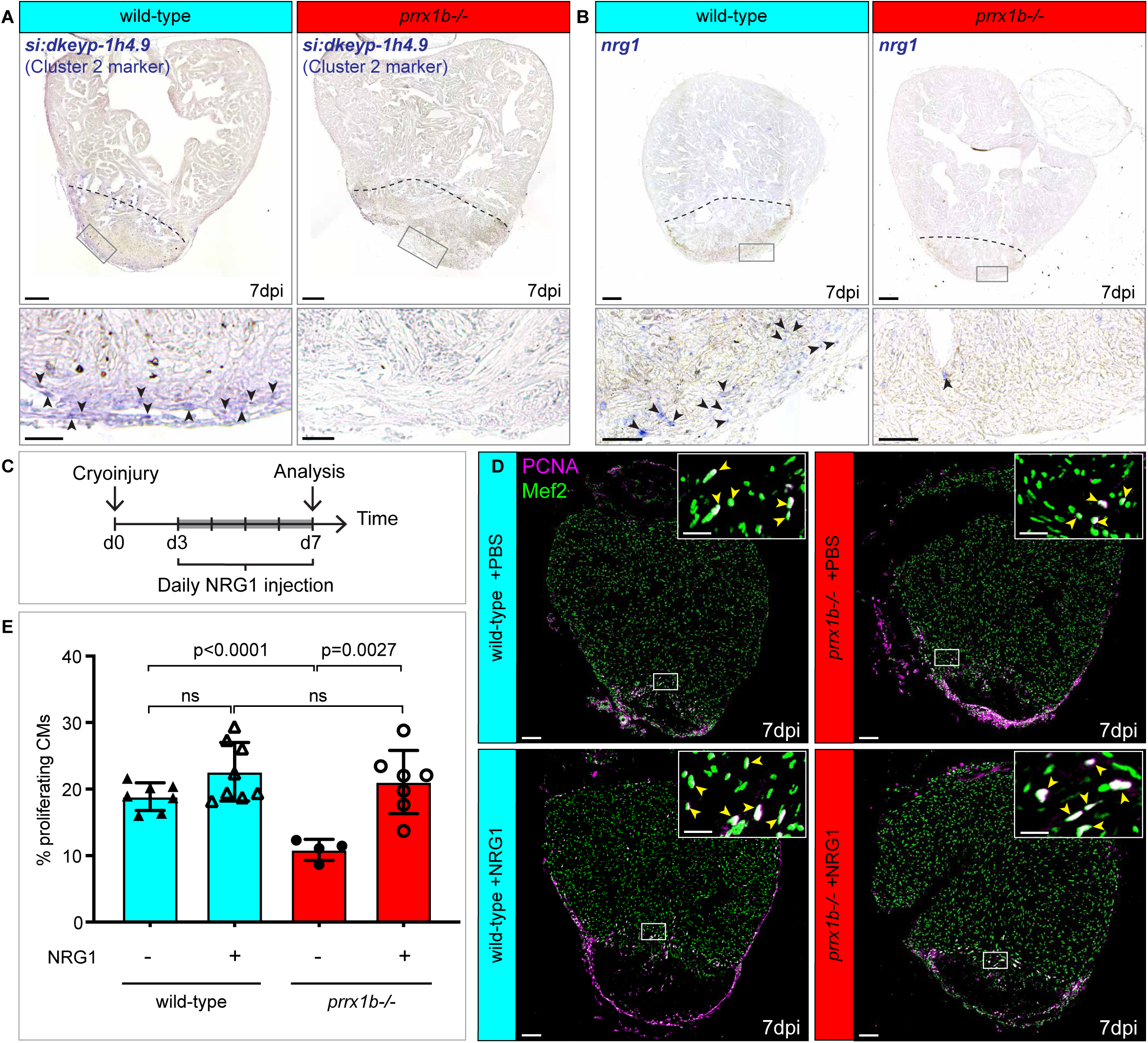
*prrx1b*-/- cardiomyocyte proliferation defect is a consequence of reduced Nrg1-expressing EPD-fibroblasts. (**A**,**B**) *In situ* hybridization for the (**A**) cluster 2 gene *si:dkeyp-1h4*.*9* or (**B**) *nrg1* in either wild-type or *prrx1b*-/- hearts at 7dpi. Lower panels show higher magnifications of boxed regions indicated in upper panels. Arrowheads point to cells with high expression. Scale bars represent 100μm in the overview images and 25μm in the zoom-ins. Hearts analysed per condition: 3 (in **A**) or 6 (in **B**). (**C**) Schematic representation of the workflow used for the NRG1 injection experiments. (**D**) Immunofluorescent staining with anti-PCNA (proliferation) and anti-Mef2 (cardiomyocytes) antibodies on 7dpi wild-type and *prrx1b*-/- heart sections injected with PBS (control) or recombinant NRG1. Arrowheads in zoom-ins indicate proliferating cardiomyocytes. Scale bars represent 100μm in the overview image and 25μm in the zoom-ins. (**E**) Quantification of the percentage of (PCNA+) proliferating border zone cardiomyocytes. ns = not significant. Error bars represent SD.

**Figure 6:**
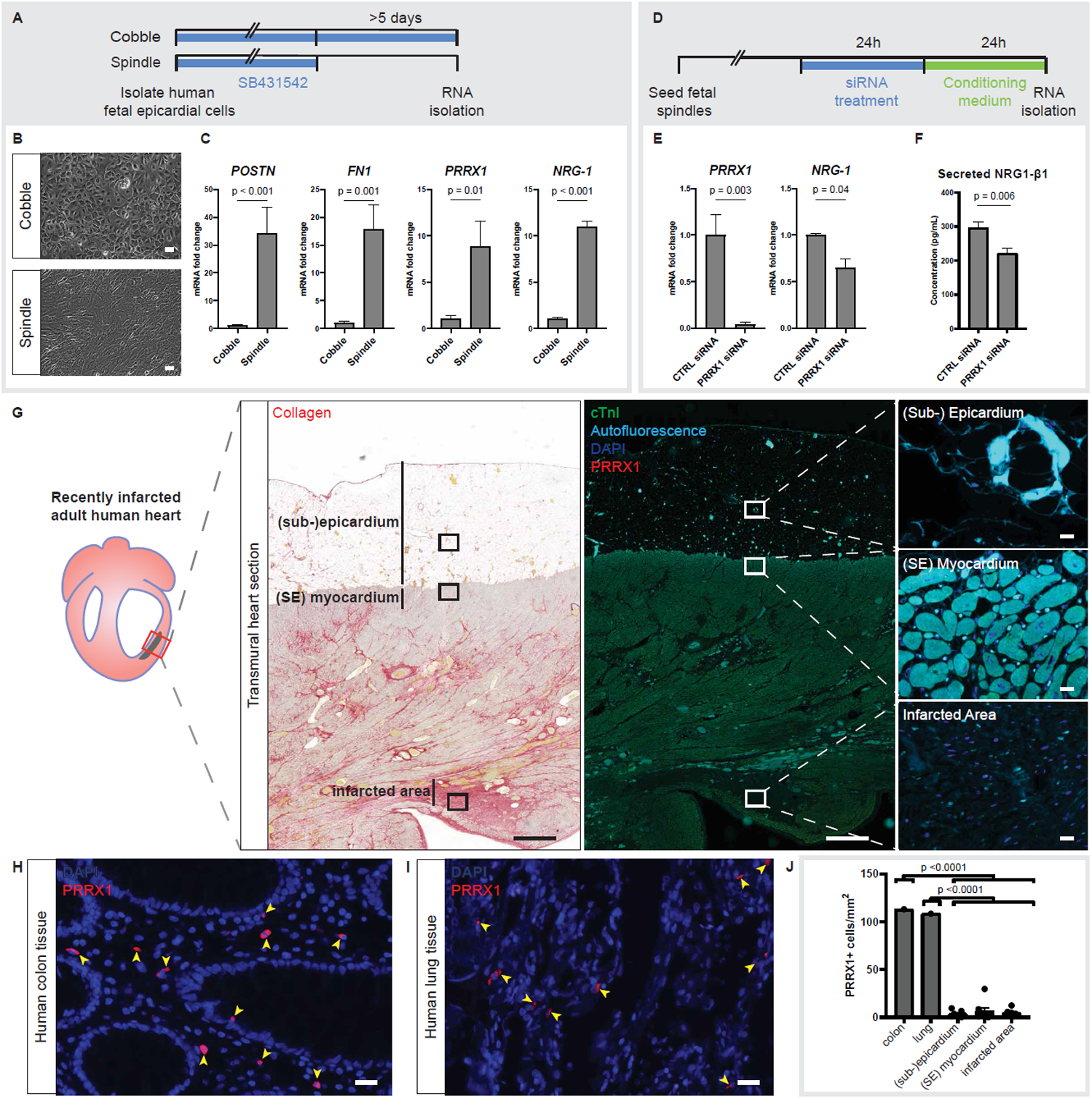
PRRX1 regulates NRG-1 secretion in human fetal epicardial cells, but is nearly undetectable in adult injured human hearts. (**A**) Schematic representation of the workflow for panels B and C. After isolation, human fetal epicardial cells are cultured in the presence of ALK4/5/7 kinase inhibitor SB431542. Cells transform from cobble- to spindle-shape upon removal of SB431542. (**B**) Representative brightfield pictures of cobble- and spindle-shaped human fetal epicardial cells. Scale bars represent 100 µm. (**C**) qPCR results for POSTN, FN1, PRRX1 and NRG1 in human fetal cobble and spindle epicardial cells (n=3-4). (**D**) Schematic representation of the workflow for panel E. (**E**) qPCR results for PRRX1 and NRG1 in human fetal spindle epicardial cells after PRRX1 siRNA treatment (n=4). (**F**) ELISA results for secreted NRG1-β1 in the conditioned cell culture medium of human fetal spindle epicardial cells between 24-48 hours after PRRX1 siRNA treatment (n=3). (**G**) Sirius red and immunofluorescent staining on consecutive human tissue samples of patients with a recent myocardial infarction for PRRX1 (red), cTnI (green) as marker for the myocardium and autofluorescence (light blue) to indicate the extracellular matrix and DAPI (dark blue). Scale bars represent 1000 µm in overview pictures and 20 µm in zoom-ins. (**H-I**) Positive control for PRRX1 staining in human colon and lung tissue. Arrowheads indicate PRRX1+ cells. Scale bars represent 20 µm. (**J**) Quantification of PRRX1+ cells in human tissue at different locations. Error bars represent SEM. For E and F, a paired t-test was used to determine statistical significance. For J, a one-way ANOVA with Dunnett’s multiple comparisons test was used to determine statistical significance. SE = sub-epicardial.

**Figure 7:**
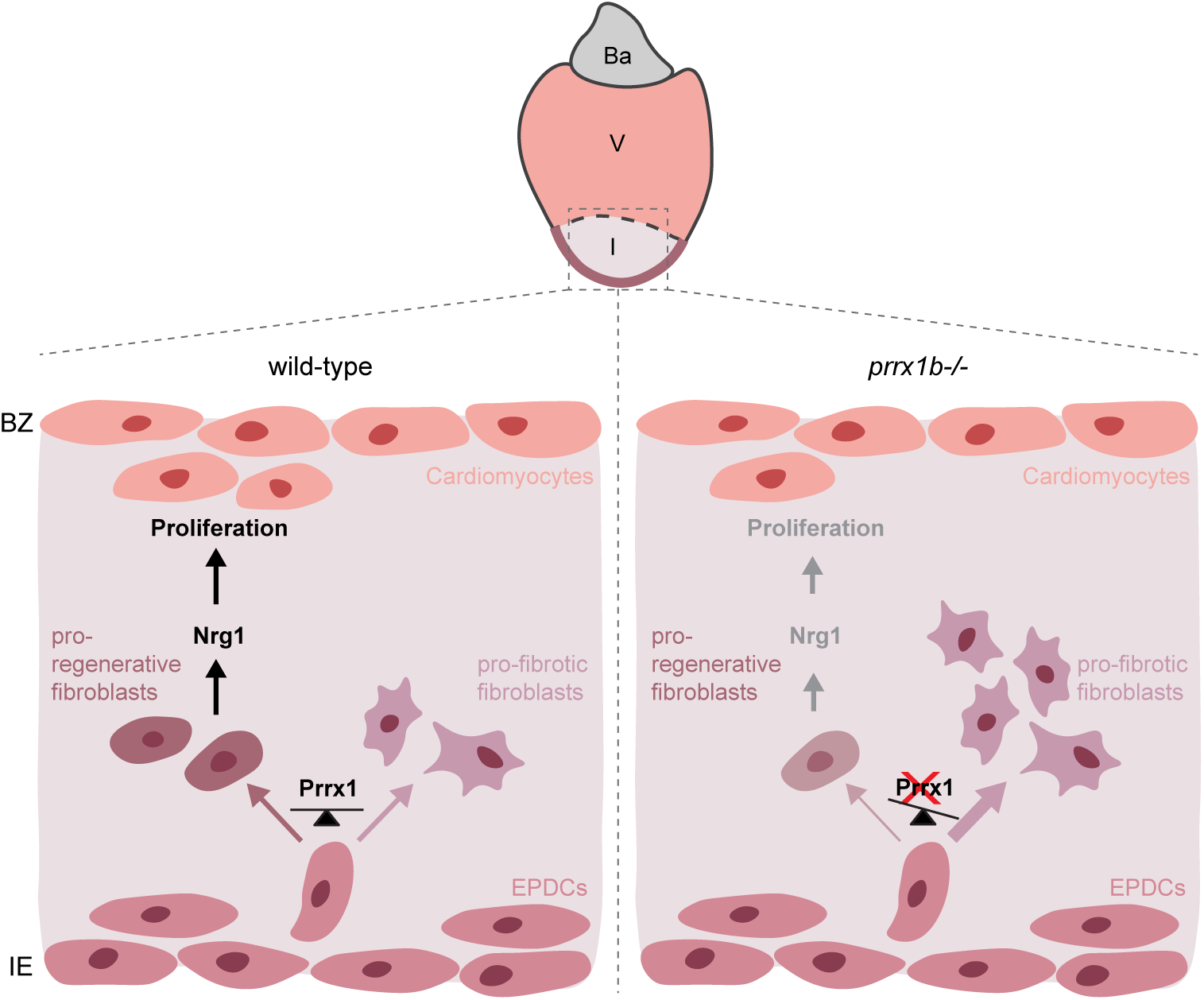
Proposed model for Prrx1b function during zebrafish heart regeneration. (Left) In a wild-type situation, the epicardium of an injured zebrafish heart gives rise to EPDCs that under influence of Prrx1b maintain a balance between pro-fibrotic (tgfb/collagen producing) fibroblasts and pro-regenerative (Nrg1 secreting) fibroblasts, allowing for regeneration. (Right) In an injured heart of *prrx1b*-/- fish, Prrx1b is no longer able to maintain this balance and EPDCs differentiate excessively in pro-fibrotic fibroblasts at the expense of pro-regenerative fibroblasts. This results in excessive scarring and reduced levels of Nrg1 leading to reduced BZ cardiomyocyte proliferation, thereby inhibiting heart regeneration. Ba = Bulbous Arteriosus; V = Ventricle; I = Injury

Taken together, these findings indicate that both *PRRX1* and *NRG1* expression is induced in human fetal EPDCs when differentiating towards a mesenchymal cell fate and that the efficient induction of NRG1 is dependent on PRRX1. This suggests that the role of PRRX1 in EPDCs is conserved between zebrafish and human. Furthermore, the near absence of PRRX1 in injured adult human hearts could party explain their limited regenerative capacity.

## Discussion

The results described here demonstrate that *prrx1b* is required for the scar-free regeneration of the injured zebrafish heart. The zebrafish genome contains a *prrx1a* and a *prrx1b* gene, which are likely the result of an ancient genome duplication that occurred in teleost (Howe et al. 2013). Our results demonstrate that while *prrx1b* is required for heart regeneration, *prrx1a* is dispensable which suggests these paralogs have non-redundant roles. This is different from their role during cartilage formation in the embryo where *prrx1a* and *prrx1b* do play redundant roles (Barske et al. 2016).

Prrx1 expression is rapidly induced in the epicardium upon injury. This is reminiscent of the induction of other genes in the epicardium such as *tbx18* and *raldh2* and implies that Prrx1 induction is part of the early activation that occurs in the entire epicardium (Cao and Poss 2018; Lepilina et al. 2006). Importantly, not all *tcf21*+ epicardial and EPDCs express Prrx1, which confirms other observations that the epicardium is a very heterogenous cell population (Cao et al. 2016; Weinberger et al. 2020).

It has been well established that EPDCs differentiate into various cell types (reviewed in Cao 2018). Retroviral labelling and Cre-mediated recombination studies in chick, mouse and zebrafish demonstrate that EPDCs differentiate into fibroblasts and vascular support cells (e.g. pericytes) (Acharya et al. 2012; Gittenberger-de Groot et al. 1998; González-Rosa et al. 2012; Kikuchi et al. 2011; Männer 1999; Mikawa and Gourdie 1996), which is in good agreement with our scRNAseq data.

There are also numerous reported observations suggesting that EPDCs can differentiate into endothelial cells and cardiomyocytes (Cai et al. 2008; Guadix et al. 2006; Katz et al. 2012; Männer 1999; Mikawa and Gourdie 1996; Smart et al. 2011; Zangi et al. 2013), although some of these observations have been questioned by others (Christoffels et al. 2009; Rudat and Kispert 2012). In our scRNAseq analysis of EPDCs recovered from the regenerating heart we did not find a cell type representing endothelial cells or cardiomyocytes, which is in agreement with earlier observations that *tcf21*-derived EPDCs in the zebrafish do not contribute to either endothelial or myocardial cell lineages (González-Rosa et al. 2012; Kikuchi et al. 2011).

Fibroblasts form one of the main contributors to ECM deposition in response to cardiac injury and are therefore an important cell type in maintaining the balance between the fibrotic and regenerative injury response (Chablais and Jazwinska 2012; Gemberling et al. 2015; Sánchez-iranzo et al. 2018). Here, we identified two distinct fibroblast populations that we refer to as a fibroblasts in either a pro-fibrotic or a pro-regenerative cell state (**Fig.7**). Both clusters represent activated fibroblasts based on their expression of periostin (*postnb*), a secreted protein expressed by activated (myo-)fibroblasts (Kanisicak et al. 2016; Sánchez-iranzo et al. 2018). The pro-fibrotic fibroblasts (cluster 3 cells) expressed all three TGF-β ligands supporting earlier findings that these ligands are expressed in the injury area to activate a pro-fibrotic response (Chablais and Jazwinska 2012). Pro-fibrotic fibroblasts express fibronectin-1 (*fn1a*) and various collagens (Sánchez-iranzo et al. 2018), which we found to be upregulated in the cluster 3 fibroblasts corroborating their pro-fibrotic nature. In *prrx1b-/-* hearts these pro-fibrotic fibroblasts were more abundant which is in good agreement with the observed excess of collagen deposition. Whereas cardiac fibrosis is permanent in the injured mammalian heart, it is resolved in the zebrafish heart. The mechanism for this regression in the zebrafish heart is not well understood. It could be related to the observation that activated fibroblasts partially return to a quiescent state (Sánchez-iranzo et al. 2018). Our observation of more activated pro-fibrotic fibroblasts in *prrx1b-/-* hearts and slowed regression of the fibrotic region suggests that an excessive fibrotic response of this type of fibroblast is detrimental for scar regression.

The other fibroblast cell state that we identified using the scRNAseq approach was assumed by cluster 2 cells which showed only limited expression of pro-fibrotic genes. Interestingly, these cluster 2 cells are nearly absent from the non-regenerating *prrx1b-/-* heart, suggesting these cells represent fibroblasts in a more pro-regenerative state. Many factors secreted by activated fibroblasts have been implicated to affect cardiac development and regeneration, suggesting that the pro-regenerative function of fibroblasts might be accomplished through their secretory role (Barandon et al. 2003; Gibb, Lavery, and Hoppler 2013; Huang et al. 2013; Li et al. 2011b; Sánchez-iranzo et al. 2018; Wu et al. 2015). In addition, experiments co-culturing fibroblasts with cardiomyocytes show that fibroblasts can induce cardiomyocyte proliferation (Ieda et al. 2009). Here we link the presence of these pro-regenerative fibroblasts with the expression of *nrg1*. Nrg1 is a potent inducer of cardiomyocyte proliferation by activation of the ErbB2 signaling pathway (Bersell et al. 2009; D’Uva et al. 2015; Gemberling et al. 2015), and its absence in *prrx1b-/-* hearts explains the observed cardiomyocyte proliferation defect. Whether Prrx1b regulates *nrg1* expression directly or indirectly remains to be elucidated. Furthermore, our observations in human fetal EPDCs correlate well with previous observations that embryonic EPDCs secrete growth factors that promote cardiomyocyte proliferation (Chen et al. 2002; Weeke-Klimp et al. 2010). Interestingly, we observed that human fetal EPDCs, which are induced to initiate EMT *in vitro*, express NRG1 and that this is dependent on PRRX1. These findings are relevant for future strategies exploiting EPDCs and EPDC-derived growth factors to stimulate human heart regeneration. Taken together, we show that Prrx1b expression in EPDCs is indispensable for the induction of a pro-regenerative fibroblast state at the expense of a pro-fibrotic fibroblast state. In doing so, Prrx1b ensures the balance between fibrotic repair and the regeneration of lost myocardium.

## Materials and Methods

### Animal experiments

Animal care and experiments conform to the Directive 2010/63/EU of the European Parliament. All animal work was approved by either the Animal Experimental Committee of the Instantie voor Dierenwelzijn Utrecht (IvD) and were performed in compliance with the Dutch government guidelines. Zebrafish were housed under standard conditions (Aleström et al. 2019).

### Zebrafish lines

The following zebrafish lines were used: TL, prrx1a, prrx1a^el558^, prrx1b^el491^ (Barske et al. 2016), Tg(tcf21:CreERT2)(Kikuchi et al. 2011); Tg(ubi:loxP-EGFP-loxP-mCherry)(Mosimann et al. 2011).

### Cryoinjuries in zebrafish

To address experiments in a regeneration context, cardiac cryo-injuries were performed on TL and prrx1b^el491^ (with and without the Tg(tcf21:CreERT2; ubi:loxP-EGFP-loxP-mCherry)) fish of ∼4 to 18 months of age. The cryoinjuries were performed as described in (Schnabel et al., 2011), with the exception of the use of a copper filament (0.3mm) cooled in liquid nitrogen instead of dry ice. Animals were excluded from the study in case of signs of aberrant behaviour/sickness/infection, according to animal guidelines.

### Histology and enzyme histochemistry

Acid fuchsin orange G (AFOG) staining was performed on paraffin sections of zebrafish ventricles as previously described (Poss et al., 2002). Paraffin sections of 7, 30 and 90dpi hearts were prepared as described below (see Methods section: *In situ* hybridization). Sirius Red staining was performed on similar paraffin sections as previously described on http://www.ihcworld.com/_protocols/special_stains/sirius_red.htm, excluding the haematoxylin step.

### Immunofluorescence

Adult zebrafish ventricles were isolated and fixed in 4% PFA (4°C O/N on shaker). The next day, the hearts were washed 3× 10 minutes in 4% sucrose phosphate buffer, after which they were incubated at RT for at least 5h in 30% sucrose phosphate buffer until the hearts floated. Then, they were embedded in cryo-medium (OCT). Cryo-sectioning of the hearts was performed at 10um thickness. Primary antibodies used include anti-PCNA (Dako#M0879, 1:800), anti-Mef2c (Santa Cruz #SC313, Biorbyt#orb256682 both 1:1000), Anti-tropomyosin (Sigma #122M4822, 1:400), Living Colors anti-DsRed (Clontech #632496, 1:100), anti-RFP (Novus Biologicals #42649), anti-Prrx1 (Gift from Tenaka lab, described in (Gerber et al. 2018; Oliveira et al. 2018), 1:200). Secondary antibodies include Anti-chicken Alexa488 (Thermofisher, #A21133, 1:500), anti-rabbit Alexa555 (Thermofisher, #A21127, 1:500), anti-mouse Cy5 (Jackson ImmunoR, #118090, 1:500). Nuclei were shown by DAPI (4’,6- diamidino-2-phenylindole) staining. Images of immunofluorescence stainings are single optical planes acquired with a Leica Sp8 confocal microscope. Human tissue was used for immunofluorescent staining as described (Kruithof 2020, CVR) using the following antibodies: anti-cTnI (Hytest, #4T21, 1:1000), anti-Prrx1 (Gift from Tenaka lab, described in (Gerber et al. 2018; Oliveira et al. 2018), 1:200). As a control, sections without primary antibody were taken along. Nuclei were shown by DAPI staining. Tissue sections were imaged using Pannoramic 250 slide scanner (3DHISTECH).

### Quantitative analyses

Unless stated otherwise, three individual sections per heart have been analysed including presented data obtained through *in situ* hybridization, immunohistochemistry and Sirius Red staining. Imaris x64 V3.2.1 was used to analyse immunofluorescent images made with a Leica SP8 confocal microscope. Proliferation percentages of border zone cardiomyocytes were determined using the spots selection tool. A region of interest (200μm) consisting of the border zone was chosen and cardiomyocytes (Mef2) were selected by classifying them as 5μm diameter or bigger. Proliferating cardiomyocytes were selected by hand using the PCNA channel. To quantify the percentage of *tcf21:*mCherry+ cell invasion into the injury, the surface selection tool was used to mark the total *tcf21:*mCherry+ area around and in the injury. The total injury area plus 100μm border zone was chosen as a region of interest. The measurement we used was the average value of the volume. Then, surfaces inside the injury were selected manually to create a subset of the total surface. Proliferation of *tcf21:*mCherry+ cells was measured as the amount of PCNA+ cells per μm^2^ of *tcf21:*mCherry+ tissue surface, since the cytoplasmic mCherry signal does not allow for the distinction between individual cells. *tcf21:*RFP+ PCNA+ cells were counted manually. ImageJ was used to quantify the remaining scar size of 30 and 90dpi heart sections following AFOG staining. All heart sections were stained, imaged and quantified for scar tissue area using ImageJ. Sirius Red staining in wild-type and *prrx1b*-/- hearts was analysed using the image-J macros MRI fibrosis tool (http://dev.mri.cnrs.fr/projects/imagej-macros/wiki/Fibrosis_Tool). Caseviewer (3DHISTECH) was used to analyse PRRX1+ cells in human cardiac tissue. Positive cells were manually counted in 10 areas of 0.1 mm^2^ in the (sub-) epicardial layer, sub-epicardial myocardium and infarcted area, that were blindly selected. As a technical positive control, 10 areas of 0.1 mm^2^ were similarly quantified in human colon and lung tissue.

### Lineage tracing of zebrafish epicardial cells

To lineage trace epicardial and epicardial derived cells, we combined the Tg(tcf21:CreERT2) with the tg(ubi:loxP-EGFP-loxP-mCherry). Both wild-type and *prrx1b*-/- embryos with a single copy of both transgenes Tg(tcf21:CreERT2; ubi:loxP-EGFP-loxP-mCherry) were incubated in 4-hydroxytamoxifen (4-OHT) as described in (Kikuchi et al. 2011; Mosimann et al. 2011) from 1dpf until 5dpf at a concentration of 5μM. At 5dpf, embryos were selected that were positive for epicardial mCherry signal and grown to adulthood.

### Isolation of single cells from cryoinjured hearts

Cryoinjured hearts of either *prrx1b* wild-type siblings (n = 20) or *prrx1b* homozygous mutants (n = 20) previously recombined as embryo (Tg(tcf21:CreERT2; ubi:loxP-EGFP-loxP-mCherry) were extracted at seven dpi. Cells were dissociated according to (Tessadori et al. 2012). For cell sorting, viable cells were gated by negative DAPI staining and positive YFP-fluorescence. In brief, the FACS gating was adjusted to sort cells positive for mCherry (recombined epicardial derived cells) and negative for EGFP (unrecombined cells). In total 1536 cells (768 *prrx1b* wild-type sibling cells and 768 *prrx1b* homozygous mutant cells) were sorted into 384-well plates and processed for mRNA sequencing as described below.

### Single-cell mRNA sequencing

Single-cell sequencing libraries were prepared using SORT-seq (Muraro et al. 2016). Live cells were sorted into 384-well plates with Vapor-Lock oil containing a droplet with barcoded primers, spike-in RNA and dNTPs, followed by heat-induced cell lysis and cDNA syntheses using a robotic liquid handler. Primers consisted of a 24 bp polyT stretch, a 4 bp random molecular barcode (UMI), a cell-specific 8 bp barcode, the 5’ Illumina TruSeq small RNA kit adapter and a T7 promoter. After cell-lysis for 5 min at 65°C, RT and second strand mixes were distributed with the Nanodrop II liquid handling platform (Inovadyne). After pooling all cells in one library, the aqueous phase was separated from the oil phase, followed by IVT transcription. The CEL-Seq2 protocol was used for library prep (Hashimshony et al. 2016). Illumina sequencing libraries were prepared with the TruSeq small RNA primers (Illumina) and paired-end sequenced at 75 bp read length on the Illumina NextSeq platform. Mapping was performed against the zebrafish reference assembly version 9 (Zv9).

### Bioinformatic analysis

To analyze the single-cell RNA-seq data, we used an updated version (RaceID3) of the previously published RaceID algorithm (Grün et al. 2015). For the adult hearts we had a dataset consisting of two different libraries of 384 cells per genotype (*prrx1b* wild-type or homozygous mutants) each for a combined dataset of 768 cells, in which we detected a total of 20995 genes. We detected an average of 7022 reads per cell. Based on the distribution of the log10 total reads plotted against the frequency, we introduced a cutoff at minimally 1000 reads per cell before further analysis. This reduced the number of cells used in the analysis to 711 wild-type and 727 mutant cells. Batch-effects were analyzed and showed no plate-specific clustering of certain clusters. The StemID algorithm were used as previously published (Grün et al. 2016). In short, StemID is an approach developed for inferring the existence of stem cell populations from single-cell transcriptomics data. StemID calculates all pairwise cell-to-cell distances (1 – Pearson correlation) and uses this to cluster similar cells into clusters that correspond to the cell types present in the tissue. The StemID algorithm calculates the number of links between clusters. This is based on the assumption that cell types with less links are more canalized while cell types with a higher number of links have a higher diversity of cell states. Besides the number of links, the StemID algorithm also calculates the change in transcriptome entropy. Differentiated cells usually express a small number of genes at high levels in order to perform cell specific functions, which is reflected by a low entropy. Stem cells and progenitor cells display a more diverse transcriptome reflected by high entropy (Banerji et al. 2013). By calculating the number of links of one cluster to other clusters and multiplying this with the change in entropy, it generates a StemID score, which is representative to ‘stemness’ of a cell population. Differential gene expression analysis was performed using the ‘diffexpnb’, which makes use of the DESeq algorithm. P-values were Benjamini-Hochberg corrected for false discovery rate to make the cutoff.

### Statistical analysis of data

Unless stated otherwise, all statistical testing was performed by unpaired T-tests.

### *In situ* hybridization

*In situ* hybridization was performed on paraffin sections. After o/n fixation in 4% PFA, hearts were washed in PBS twice, dehydrated in EtOH, and embedded in paraffin. Serial sections were made at 8um thickness. *In situ* hybridization was performed as previously described (Moorman et al., 2001) except that the hybridization buffer used did not contain heparin and yeast total RNA. When *in situ* hybridization was done for multiple probes, INT-BCIP staining solution (red/brown staining) was used for the additional probe instead of NBT-BCIP (blue staining).

### Intraperitoneal injections in zebrafish

Intraperitoneal injections of human recombinant NRG1 (peptrotech: recombinant human heregulin-b1, catalog#:100-03) were performed as described by Kinkel et al. (Kinkel et al. 2010). Fish were sedated using MS222 (0.032% wt/vol). Injections were performed using a Hamilton syringe (Gauge 30), cleaned before use by washing in 70% ethanol followed by 2 washes in PBS. Injection volumes were adjusted on the weight of the fish (30ul/g) and a single injection contained 60 ug/kg (diluted in PBS/BSA 0.1%).

### Human epicardial cell culture

Human fetal embryonic hearts of a gestational age between 12-18 weeks were collected anonymously and under informed consent from abortion material after elective abortion. Epicardial cells were isolated as described in (Dronkers et al. 2018). Cells were cultured in Dulbecco’s modified Eagle’s medium (DMEM low-glucose, Gibco) and Medium 199 (M199, Gibco) mixed in a 1:1 ratio, supplemented with 10% fetal bovine serum (heat inactivated for 25 minutes at 56 °C, Gibco), 100 U/mL penicillin (Roth), 100 mg/mL streptomycin (Roth), and 10 μM ALK4/5/7 kinase inhibitor SB431542 (Tocris) at 37 °C in 5% CO2. EMT was induced by removal of SB431542 from the medium. This research was carried out according to the official guidelines of the Leiden University Medical Center and approved by the local Medical Ethics Committee. This research conforms to the Declaration of Helsinki.

### PRRX1 KD in human epicardial cells

Human fetal epicardial cells were treated with SMARTpool ON-TARGETplus PRRX1 or a non-targeting control siRNA according to the manufacturers’ protocol in a concentration of 25 nM (Dharmacon). After 48 hours, cells were collected for qPCR. All experiments were performed with three or four individual cell isolations.

### Human material used for immunohistochemistry

Healthy lung and colon tissue was obtained from the normal tissue biobank of the University Medical Center Utrecht and was fixed in PFA and embedded in paraffin. Paraffin-embedded infarcted human heart tissue from 8 individuals who had recently died of MI were retrieved from the pathology archive of the University Medical Center Utrecht. Material was handled in a coded manner that met the criteria of the Code of Conduct used in the Netherlands for the responsible use of human tissue in medical research (www.federa.org/codes-conduct). Collection of the archive material was approved by the local biobank review committee (protocol 15–252).

### qPCR

ReliaPrep™ RNA Miniprep Systems (Promega) was used to isolate mRNA, of which the concentration and purity were determined using NanoDrop 1000 Spectrophotometer (Thermo Fisher Scientific). cDNA synthesis was performed using the RevertAid H Minus First Strand cDNA Synthesis Kit (Thermo Fisher Scientific). Next, qPCR was performed using SYBR Green (Promega) and run on a C1000 Touch™ thermal cycler (Bio-Rad). All samples were run in triplicates, expression levels were corrected for primer efficiency and normalized for two reference genes (TBP and HPRT1).

#### Primer sequences

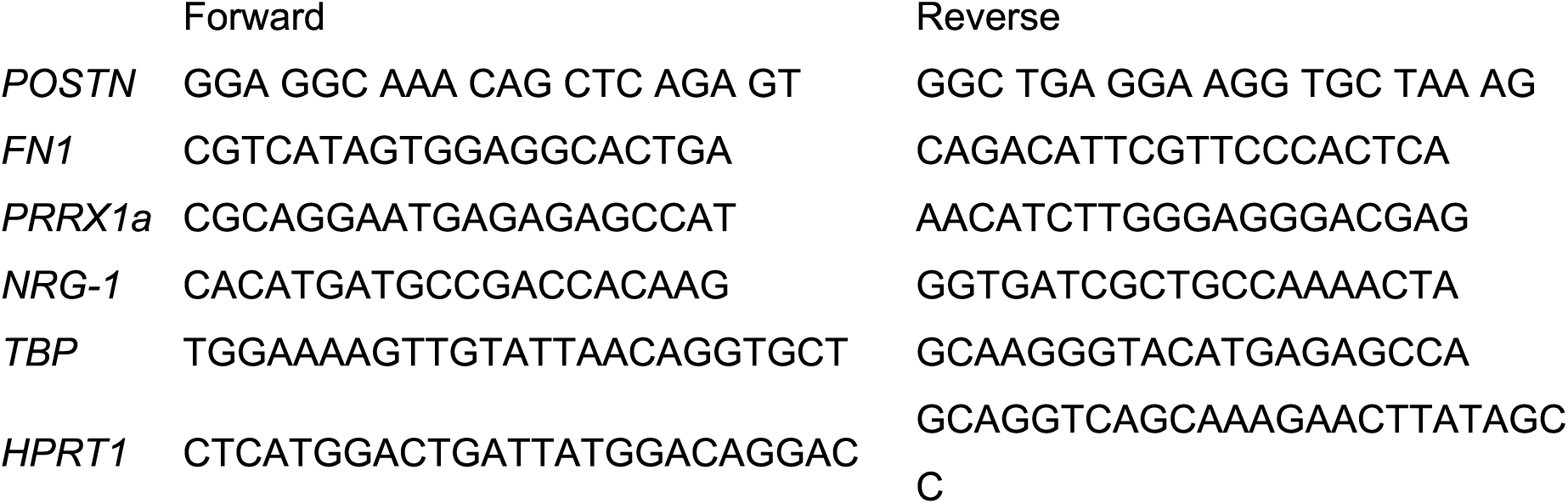

### ELISA

Conditioned medium was collected for 24 hours after 24 hours of siRNA treatment, centrifuged and immediately frozen at -20°. Cell culture medium was taken along as a control sample. An NRG1-β1 ELISA assay was performed according to the protocol of the manufacturer (Human NRG1-β1 DuoSet ELISA, R&D systems). Absolute NRG1-β1 concentration was calculated based on the standard curve.

---

**Supplementary Figure 1:**
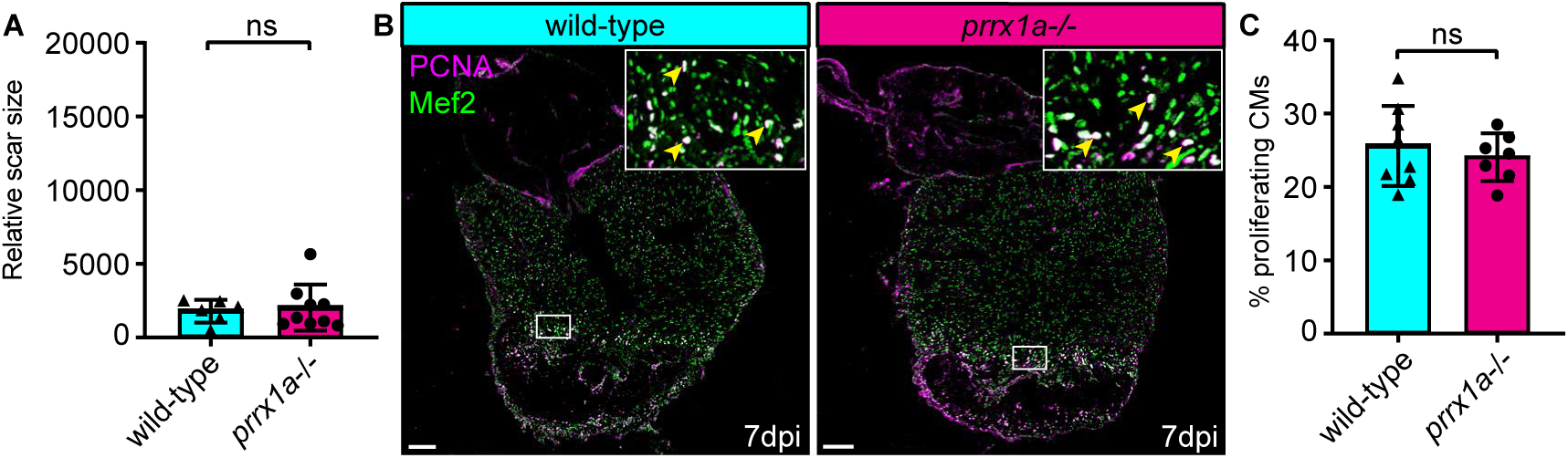
*prrx1a* is dispensable for zebrafish border zone cardiomyocyte proliferation and heart regeneration. (**A**) Quantification of the remaining scar size at 30dpi shows no significant difference between *prrx1a*-/- hearts and wild-type siblings. Error bars represent SD. (**B**) Immunofluorescent staining on 7dpi wild-type and *prrx1a*-/- heart sections using an anti-Mef2 antibody as a marker for cardiomyocyte nuclei, and an anti-PCNA antibody as a nuclear proliferation marker. Arrowheads in zoom-ins indicate proliferating cardiomyocytes. Scale bars represent 100μm in the overview images and 10μm in the zoom-ins. (**C**) Quantification of the percentage of (PCNA+) proliferating border zone cardiomyocytes shows no significant difference between *prrx1a*-/- hearts and their wild-type siblings. Error bars represent SD.

**Supplementary Figure 2:**
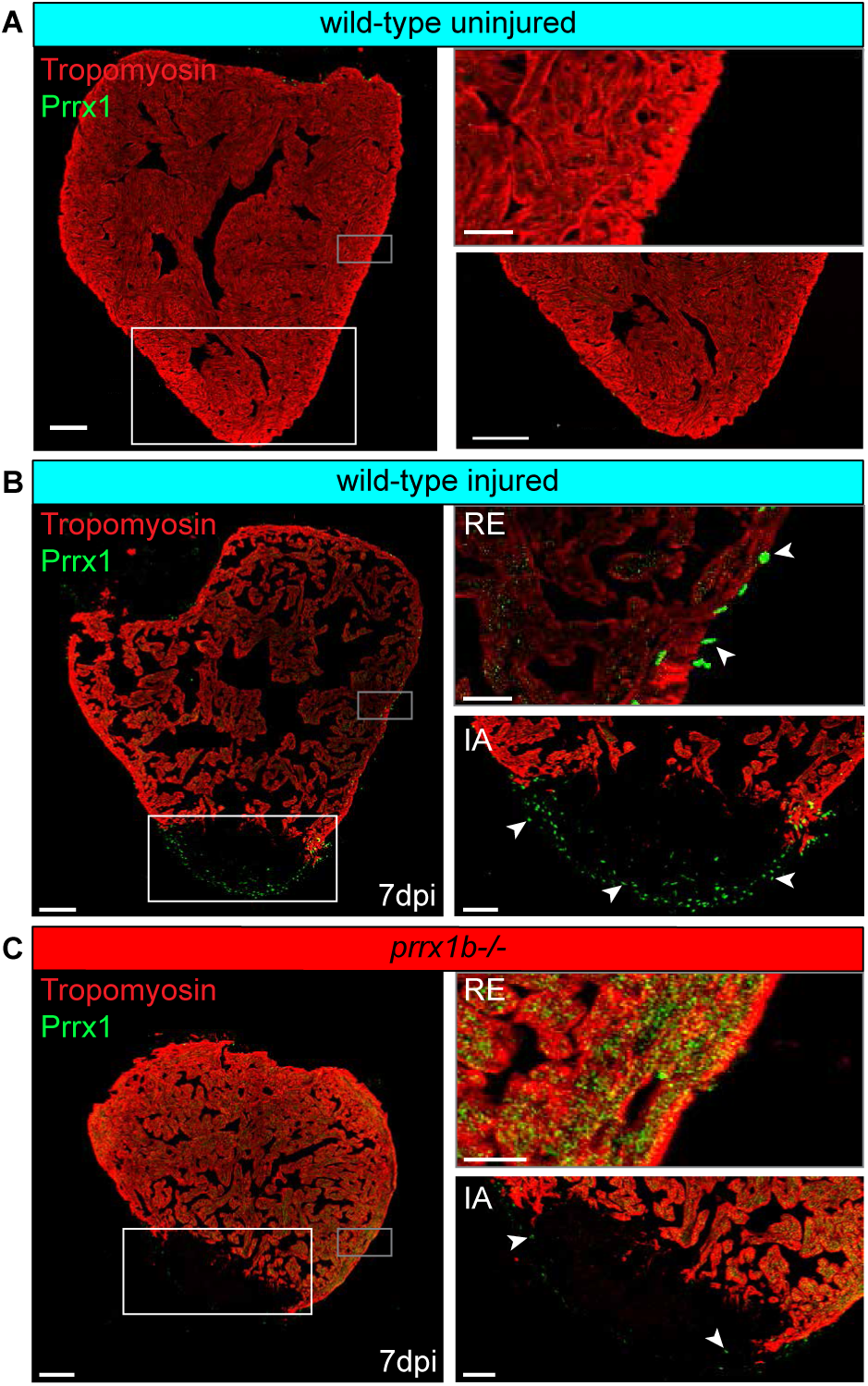
Injured zebrafish hearts express Prrx1 in cells surrounding the injury area, which is severely reduced in the *prrx1b*-/- hearts. (**A-C**) Confocal images of immunohistochemistry against tropomyosin staining CM nuclei (red) and Prrx1 protein (green) in (**A**) uninjured wildtype hearts (**B**) injured wild-type hearts and (**C**) injured *prrx1b*-/- hearts at 7 dpi. Right panels show magnifications of boxed areas displayed with arrowheads pointing to Prrx1+ cells. Note strong reduction in number of Prrx1+ cells in (**C**). Scale bars represent 100μm in the overview images, 20μm in the top zoom-ins and 50μm in the bottom zoom-ins. Hearts analyzed per condition: 3. IA = Injury Area; RE = Remote Epicardium.

**Supplementary Figure 3:**
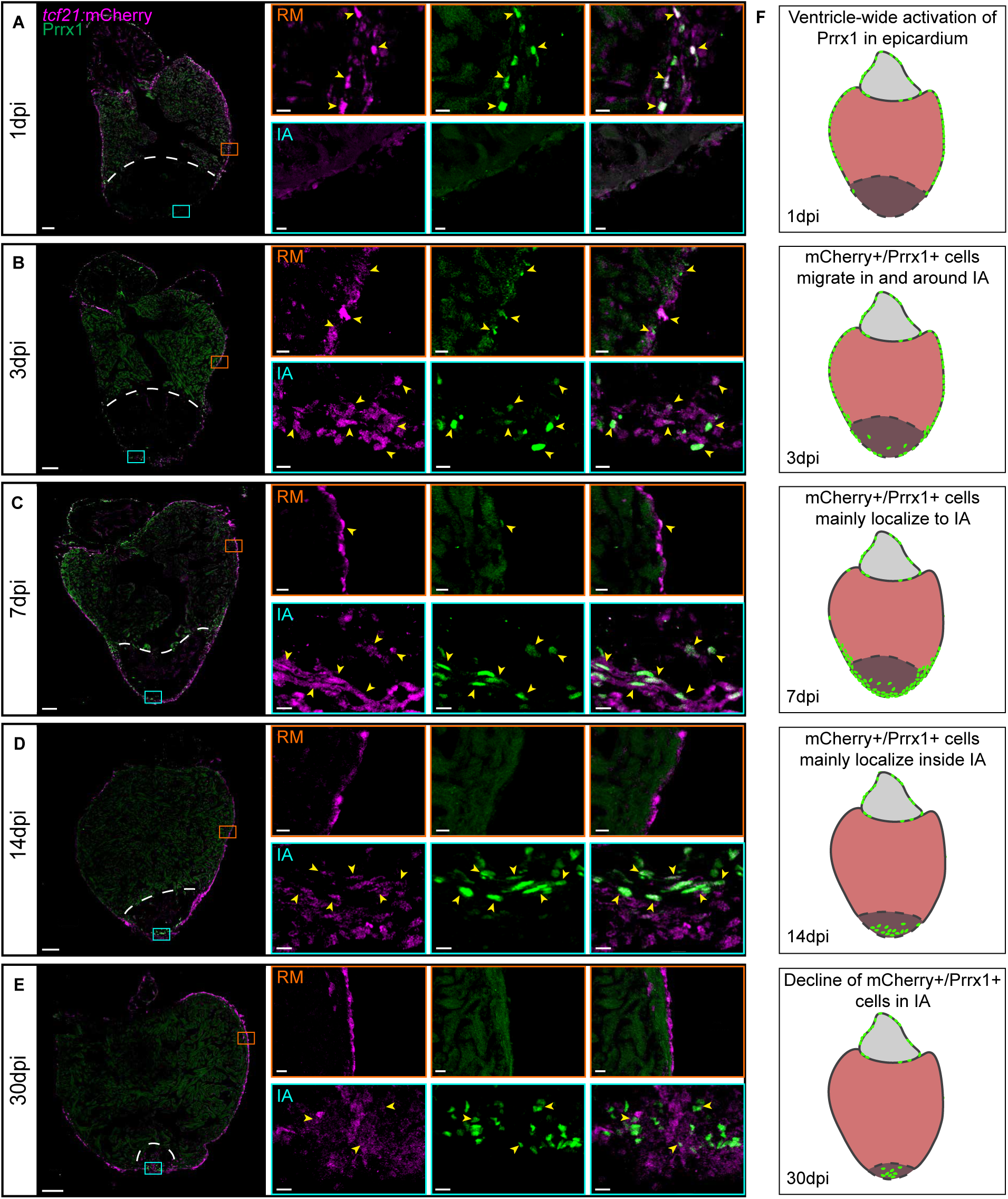
Prrx1 is expressed in EPDC and follows epicardial dynamics post injury. (**A-E**) Immunofluorescent staining on uninjured, 1, 3, 7, 14 and 30dpi wild-type heart sections. Immunofluorescent staining on tcf21:mCherry+ wild-type heart sections using an antibodies against Prrx1 (green) and mCherry (magenta) to mark tcf21:mCherry+ cells. Dotted line represents injury border. Arrowheads indicate tcf21:mCherry+/Prrx1+ cells. Scale bars represent 100μm in the overview images and 10μm in the zoom-ins. RM= Remote myocardium, IA= Injury area. Hearts analysed per condition: 6. (**F**) Schematic representation of Prrx1 dynamics upon injury. Prrx1+ cells are in green. Dark color at the apex represents the injury area.

**Supplementary Figure 4:**
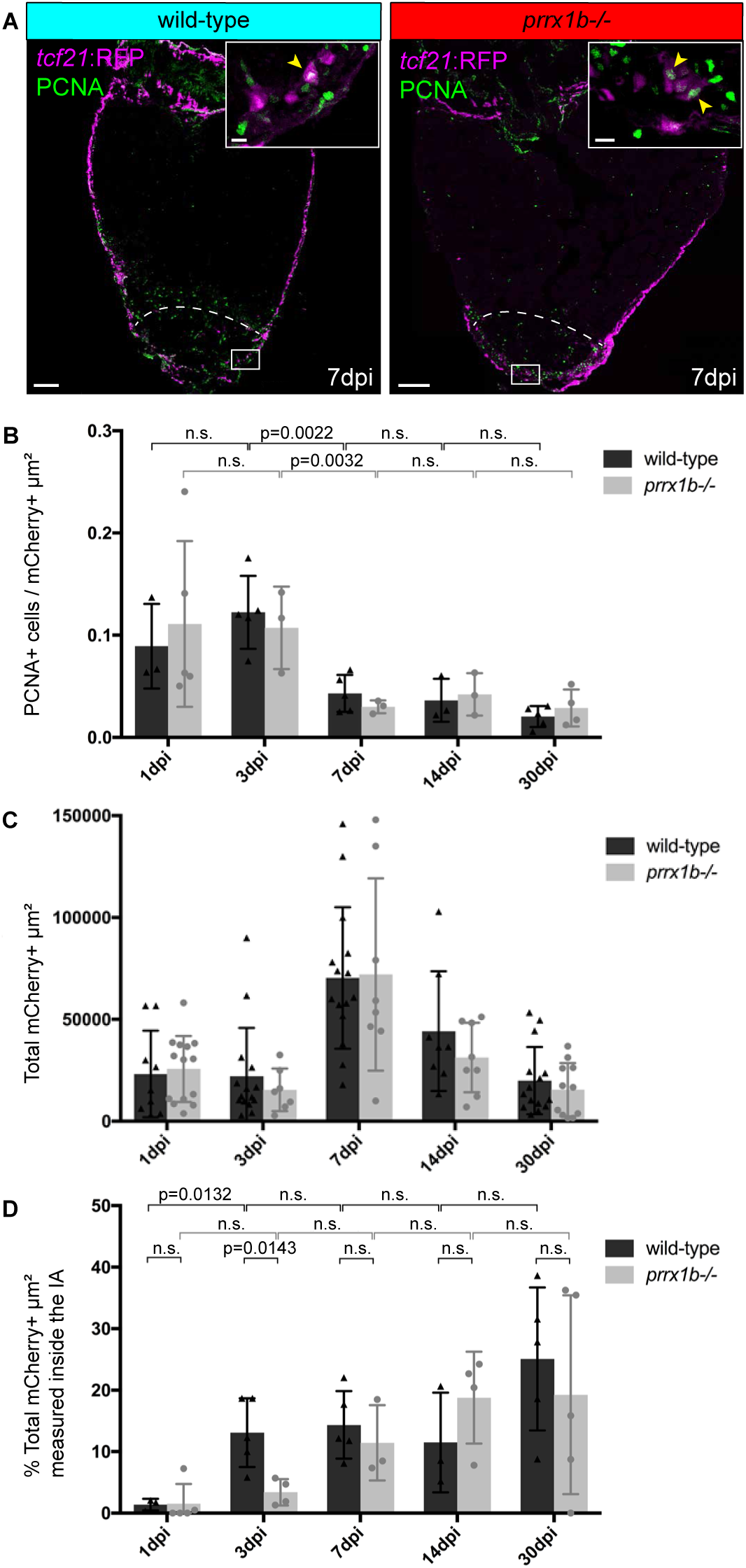
Quantification of invasion and proliferation of tcf21:mCherry+ cells at 1, 3, 7, 14 and 30dpi in wild-type and *prrx1b*-/- hearts. (**A**) Confocal images showing PCNA (green) and tcf21:mCherry (magenta) in wild-type sibling and *prrx1b*-/- hearts at 7 dpi. Scalebar = 100μm, scale bar in zoom in = 10μm. (**B**) Proliferation of tcf21:mCherry+ cells was quantified as the amount of PCNA+ cells per μ m^2^ of tcf21:mCherry+ tissue surface inside and around the injury area (IA) at 1, 3, 7, 14 and 30dpi in wild-type and *prrx1b*-/- hearts. (**C**) Total surface area mCherry+ cells in μm^2^ inside and around the IA at different time points post injury. (**D**) Percentage of the total tcf21:mCherry+ um^2^ inside and around the IA found inside the injury area at 1, 3, 7, 14 and 30dpi in wild-type and *prrx1b*-/- hearts. Error bars represent SD. n.s. = not significant.

**Supplementary Figure 5:**
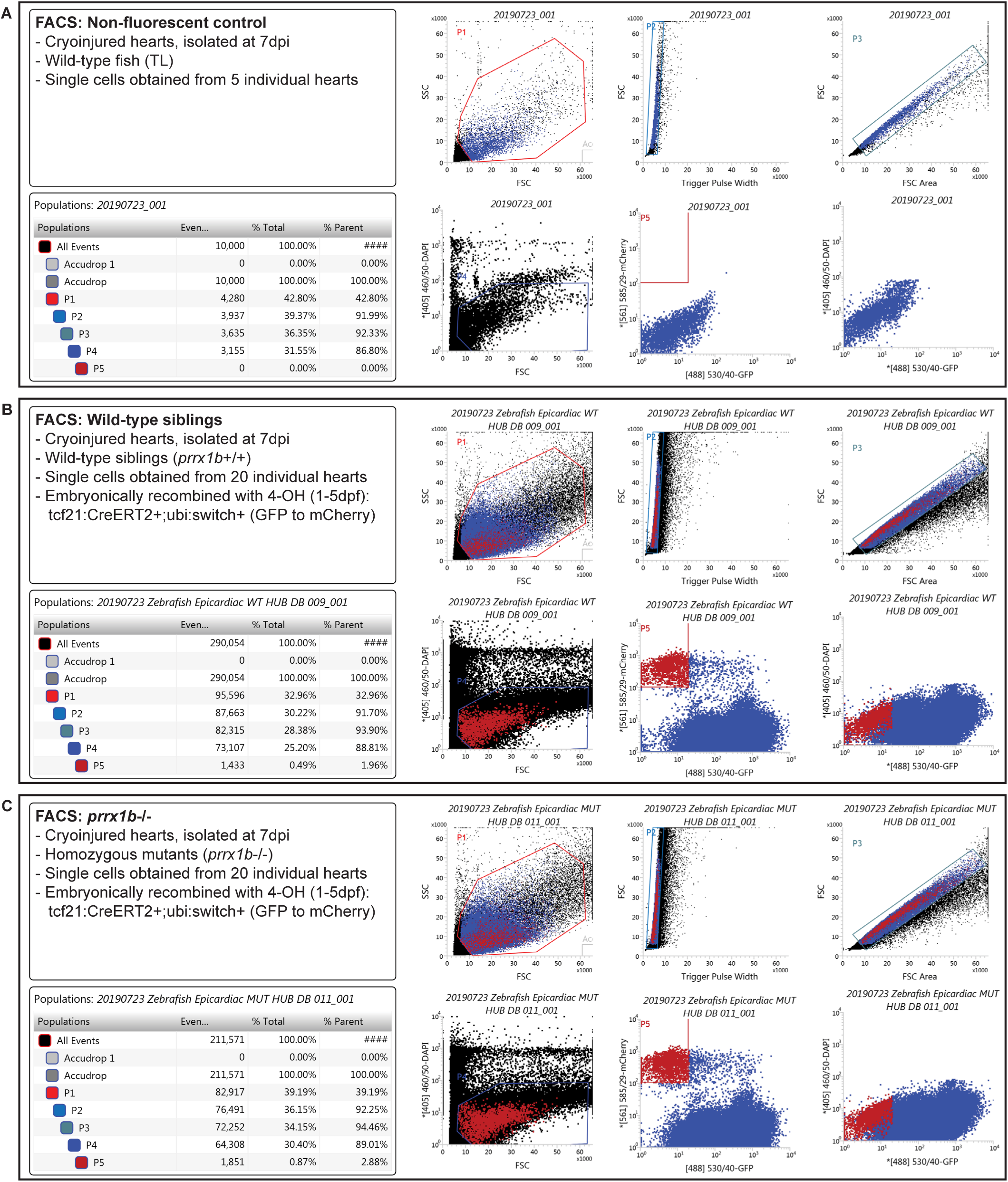
Gating details for FACS sorting. (**A-C**) Gates (P1-P5) used to sort out mCherry+/GFP-cells used for single cell sequencing analysis.

**Supplementary Figure 6:**
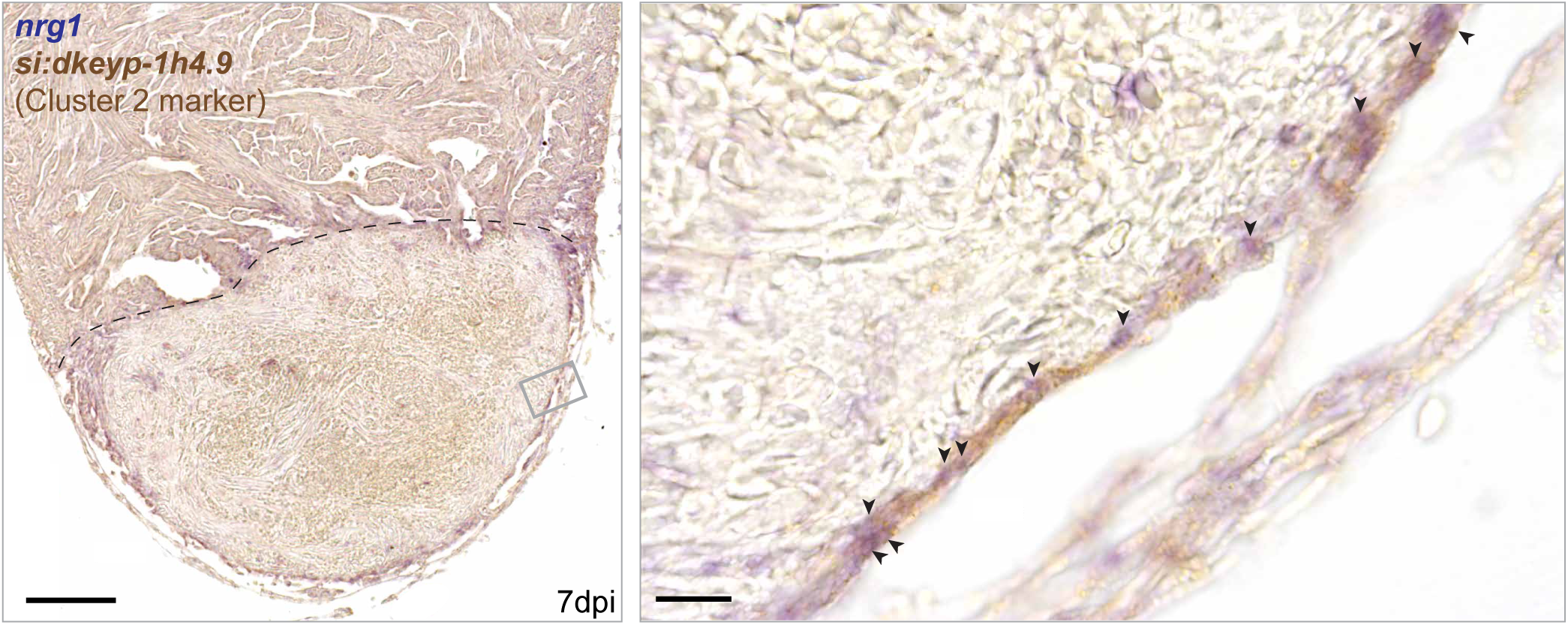
Cluster 2 fibroblasts express *nrg1*. Two-color *in situ* hybridization for *nrg1* (blue) and cluster 2 marker *si:dke-yp-1h4*.*9* (brown). Left panel shows section of injured wild-type heart at 7dpi with injury border indicated with dashed line. Right panel shows magnification of boxed region. Arrowheads indicate *nrg1* and *si:dkeyp-1h4*.*9* double labeling in epicardial region. Scale bars represent 100μm in the overview image and 10μm in the magnification. Hearts analysed per condition: 3.

## Bibliography

Acharya, Asha, Seung Tae Baek, Guo Huang, Banu Eskiocak, Sean Goetsch, Caroline Y. Sung, Serena Banfi, Marion F. Sauer, Gregory S. Olsen, Jeremy S. Duffield, Eric N. Olson, and Michelle D. Tallquist. 2012. “The BHLH Transcription Factor Tcf21 Is Required for Lineage-Specific EMT of Cardiac Fibroblast Progenitors.” Development (Cambridge, England) 139(12):2139–49.

Aleström, Peter, Livia D’Angelo, Paul J. Midtlyng, Daniel F. Schorderet, Stefan Schulte-Merker, Frederic Sohm, and Susan Warner. 2019. “Zebrafish: Housing and Husbandry Recommendations.” Laboratory Animals.

Banerji, Christopher R. S., Diego Miranda-Saavedra, Simone Severini, Martin Widschwendter, Tariq Enver, Joseph X. Zhou, and Andrew E. Teschendorff. 2013. “Cellular Network Entropy as the Energy Potential in Waddington’s Differentiation Landscape.” Scientific Reports 3.

Barandon, Laurent, Thierry Couffinhal, Jérome Ezan, Pascale Dufourcq, Pierre Costet, Philippe Alzieu, Lionel Leroux, Catherine Moreau, Danièle Dare, and Cécile Duplàa. 2003. “Reduction of Infarct Size and Prevention of Cardiac Rupture in Transgenic Mice Overexpressing FrzA.” Circulation 108(18):2282–89.

Barrallo-Gimeno, Alejandro, and M. Angela Nieto. 2005. “The Snail Genes as Inducers of Cell Movement and Survival: Implications in Development and Cancer.” Development.

Barske, Lindsey, Amjad Askary, Elizabeth Zuniga, Bartosz Balczerski, Paul Bump, James T. Nichols, and J. Gage Crump. 2016. “Competition between Jagged-Notch and Endothelin1 Signaling Selectively Restricts Cartilage Formation in the Zebrafish Upper Face.” PLoS Genetics 12(4):e1005967.

Baudino, Troy A., Wayne Carver, Wayne Giles, and Thomas K. Borg. 2006. “Cardiac Fibroblasts: Friend or Foe?” American Journal of Physiology. Heart and Circulatory Physiology 291(3):H1015–26.

Bersell, Kevin, Shima Arab, Bernhard Haring, and Bernhard Kühn. 2009. “Neuregulin1/ErbB4 Signaling Induces Cardiomyocyte Proliferation and Repair of Heart Injury.” Cell.

Cai, Chen Leng, Jody C. Martin, Yunfu Sun, Li Cui, Lianchun Wang, Kunfu Ouyang, Lei Yang, Lei Bu, Xingqun Liang, Xiaoxue Zhang, William B. Stallcup, Christopher P. Denton, Andrew McCulloch, Ju Chen, and Sylvia M. Evans. 2008. “A Myocardial Lineage Derives from Tbx18 Epicardial Cells.” Nature.

Cao, Jingli, Adam Navis, Ben D. Cox, Amy L. Dickson, Matthew Gemberling, Ravi Karra, Michel Bagnat, and Kenneth D. Poss. 2015. “Single Epicardial Cell Transcriptome Sequencing Identifies Caveolin-1 as an Essential Factor in Zebrafish Heart Regeneration.” Development (Cambridge, England) (December).

Cao, Jingli, Adam Navis, Ben D. Cox, Amy L. Dickson, Matthew Gemberling, Ravi Karra, Michel Bagnat, and Kenneth D. Poss. 2016. “Single Epicardial Cell Transcriptome Sequencing Identifies Caveolin 1 as an Essential Factor in Zebrafish Heart Regeneration.” Development (Cambridge) 143(2):232–43.

Cao, Jingli, and Kenneth D. Poss. 2018. “The Epicardium as a Hub for Heart Regeneration.” Nature Reviews Cardiology.

Chablais, Fabian, and Anna Jazwinska. 2012. “The Regenerative Capacity of the Zebrafish Heart Is Dependent on TGFβ Signaling.” Development (Cambridge, England) 139(11):1921–30.

Chablais, Fabian, Julia Veit, Gregor Rainer, and Anna Jazwinska. 2011. “The Zebrafish Heart Regenerates after Cryoinjury-Induced Myocardial Infarction.” BMC Developmental Biology 11:21.

Chen, Tim H. P., Tsai Ching Chang, Ji One Kang, Bibha Choudhary, Takako Makita, Chanh M. Tran, John B. E. Burch, Hoda Eid, and Henry M. Sucov. 2002. “Epicardial Induction of Fetal Cardiomyocyte Proliferation via a Retinoic Acid-Inducible Trophic Factor.” Developmental Biology.

Christoffels, Vincent M., Thomas Grieskamp, Julia Norden, Mathilda T. M. Mommersteeg, Carsten Rudat, and Andreas Kispert. 2009. “Tbx18 and the Fate of Epicardial Progenitors.” Nature.

D’Uva, Gabriele, Alla Aharonov, Mattia Lauriola, David Kain, Yfat Yahalom-Ronen, Silvia Carvalho, Karen Weisinger, Elad Bassat, Dana Rajchman, Oren Yifa, Marina Lysenko, Tal Konfino, Julius Hegesh, Ori Brenner, Michal Neeman, Yosef Yarden, Jonathan Leor, Rachel Sarig, Richard P. Harvey, and Eldad Tzahor. 2015. “ERBB2 Triggers Mammalian Heart Regeneration by Promoting Cardiomyocyte Dedifferentiation and Proliferation.” Nature Cell Biology 17(5).

Dronkers, Esther, Asja T. Moerkamp, Tessa van Herwaarden, Marie José Goumans, and Anke M. Smits. 2018. “The Isolation and Culture of Primary Epicardial Cells Derived from Human Adult and Fetal Heart Specimens.” Journal of Visualized Experiments.

Gemberling, Matthew, Ravi Karra, Amy L. Dickson, and Kenneth D. Poss. 2015. “Nrg1 Is an Injury-Induced Cardiomyocyte Mitogen for the Endogenous Heart Regeneration Program in Zebrafish.” ELife 4:1–17.

Gerber, Tobias, Prayag Murawala, Dunja Knapp, Wouter Masselink, Maritta Schuez, Sarah Hermann, Malgorzata Gac-Santel, Sergej Nowoshilow, Jorge Kageyama, Shahryar Khattak, Joshua D. Currie, J. Gray Camp, Elly M. Tanaka, and Barbara Treutlein. 2018. “Single-Cell Analysis Uncovers Convergence of Cell Identities during Axolotl Limb Regeneration.” Science 362(6413).

Gibb, Natalie, Danielle L. Lavery, and Stefan Hoppler. 2013. “Sfrp1 Promotes Cardiomyocyte Differentiation in Xenopus via Negative-Feedback Regulation of Wnt Signaling.” Development (Cambridge) 140(7):1537–49.

Gittenberger-de Groot, A. C., M. P. Vrancken Peeters, M. M. Mentink, R. G. Gourdie, and R. E. Poelmann. 1998. “Epicardium-Derived Cells Contribute a Novel Population to the Myocardial Wall and the Atrioventricular Cushions.” Circulation Research 82(10):1043–52.

Gittenberger-De Groot, Adriana C., Mark Paul F. M. Vrancken Peeters, Maarten Bergwerff, Monica M. T. Mentink, and Robert E. Poelmann. 2000. “Epicardial Outgrowth Inhibition Leads to Compensatory Mesothelial Outflow Tract Collar and Abnormal Cardiac Septation and Coronary Formation.” Circulation Research.

González-Rosa, Juan Manuel, Víctor Martín, Marina Peralta, Miguel Torres, and Nadia Mercader. 2011. “Extensive Scar Formation and Regression during Heart Regeneration after Cryoinjury in Zebrafish.” Development (Cambridge, England) 138(9):1663–74.

González-Rosa, Juan Manuel, Marina Peralta, and Nadia Mercader. 2012. “Pan-Epicardial Lineage Tracing Reveals That Epicardium Derived Cells Give Rise to Myofibroblasts and Perivascular Cells during Zebrafish Heart Regeneration.” Developmental Biology 370(2):173–86.

Grün, Dominic, Anna Lyubimova, Lennart Kester, Kay Wiebrands, Onur Basak, Nobuo Sasaki, Hans Clevers, and Alexander van Oudenaarden. 2015. “Single-Cell Messenger RNA Sequencing Reveals Rare Intestinal Cell Types.” Nature 525(7568):251–55.

Grün, Dominic, Mauro J. Muraro, Jean Charles Boisset, Kay Wiebrands, Anna Lyubimova, Gitanjali Dharmadhikari, Maaike van den Born, Johan van Es, Erik Jansen, Hans Clevers, Eelco J. P. de Koning, and Alexander van Oudenaarden. 2016. “De Novo Prediction of Stem Cell Identity Using Single-Cell Transcriptome Data.” Cell Stem Cell 19(2):266–77.

Guadix, Juan A., Rita Carmona, Ramón Muñoz-Chápuli, and José M. Pérez-Pomares. 2006. “In Vivo and in Vitro Analysis of the Vasculogenic Potential of Avian Proepicardial and Epicardial Cells.” Developmental Dynamics.

Guichet, Pierre Olivier, Sophie Guelfi, Marisa Teigell, Liesa Hoppe, Norbert Bakalara, Luc Bauchet, Hugues Duffau, Katrin Lamszus, Bernard Rothhut, and Jean Philippe Hugnot. 2015. “Notch1 Stimulation Induces a Vascularization Switch with Pericyte-like Cell Differentiation of Glioblastoma Stem Cells.” Stem Cells.

Hashimshony, Tamar, Naftalie Senderovich, Gal Avital, Agnes Klochendler, Yaron de Leeuw, Leon Anavy, Dave Gennert, Shuqiang Li, Kenneth J. Livak, Orit Rozenblatt-Rosen, Yuval Dor, Aviv Regev, and Itai Yanai. 2016. “CEL-Seq2: Sensitive Highly-Multiplexed Single-Cell RNA-Seq.” Genome Biology 17(1).

Howe, Kerstin, Matthew D. Clark, Carlos F. Torroja, James Torrance, Camille Berthelot, Matthieu Muffato, John E. Collins, Sean Humphray, Karen McLaren, Lucy Matthews, Stuart McLaren, Ian Sealy, Mario Caccamo, Carol Churcher, Carol Scott, Jeffrey C. Barrett, Romke Koch, Gerd Jörg Rauch, Simon White, William Chow, Britt Kilian, Leonor T. Quintais, José A. Guerra-Assunção, Yi Zhou, Yong Gu, Jennifer Yen, Jan Hinnerk Vogel, Tina Eyre, Seth Redmond, Ruby Banerjee, Jianxiang Chi, Beiyuan Fu, Elizabeth Langley, Sean F. Maguire, Gavin K. Laird, David Lloyd, Emma Kenyon, Sarah Donaldson, Harminder Sehra, Jeff Almeida-King, Jane Loveland, Stephen Trevanion, Matt Jones, Mike Quail, Dave Willey, Adrienne Hunt, John Burton, Sarah Sims, Kirsten McLay, Bob Plumb, Joy Davis, Chris Clee, Karen Oliver, Richard Clark, Clare Riddle, David Eliott, Glen Threadgold, Glenn Harden, Darren Ware, Beverly Mortimer, Giselle Kerry, Paul Heath, Benjamin Phillimore, Alan Tracey, Nicole Corby, Matthew Dunn, Christopher Johnson, Jonathan Wood, Susan Clark, Sarah Pelan, Guy Griffiths, Michelle Smith, Rebecca Glithero, Philip Howden, Nicholas Barker, Christopher Stevens, Joanna Harley, Karen Holt, Georgios Panagiotidis, Jamieson Lovell, Helen Beasley, Carl Henderson, Daria Gordon, Katherine Auger, Deborah Wright, Joanna Collins, Claire Raisen, Lauren Dyer, Kenric Leung, Lauren Robertson, Kirsty Ambridge, Daniel Leongamornlert, Sarah McGuire, Ruth Gilderthorp, Coline Griffiths, Deepa Manthravadi, Sarah Nichol, Gary Barker, Siobhan Whitehead, Michael Kay, Jacqueline Brown, Clare Murnane, Emma Gray, Matthew Humphries, Neil Sycamore, Darren Barker, David Saunders, Justene Wallis, Anne Babbage, Sian Hammond, Maryam Mashreghi-Mohammadi, Lucy Barr, Sancha Martin, Paul Wray, Andrew Ellington, Nicholas Matthews, Matthew Ellwood, Rebecca Woodmansey, Graham Clark, James Cooper, Anthony Tromans, Darren Grafham, Carl Skuce, Richard Pandian, Robert Andrews, Elliot Harrison, Andrew Kimberley, Jane Garnett, Nigel Fosker, Rebekah Hall, Patrick Garner, Daniel Kelly, Christine Bird, Sophie Palmer, Ines Gehring, Andrea Berger, Christopher M. Dooley, Zübeyde Ersan-Ürün, Cigdem Eser, Horst Geiger, Maria Geisler, Lena Karotki, Anette Kirn, Judith Konantz, Martina Konantz, Martina Oberländer, Silke Rudolph-Geiger, Mathias Teucke, Kazutoyo Osoegawa, Baoli Zhu, Amanda Rapp, Sara Widaa, Cordelia Langford, Fengtang Yang, Nigel P. Carter, Jennifer Harrow, Zemin Ning, Javier Herrero, Steve M. J. Searle, Anton Enright, Robert Geisler, Ronald H. A. Plasterk, Charles Lee, Monte Westerfield, Pieter J. De Jong, Leonard I. Zon, John H. Postlethwait, Christiane Nüsslein-Volhard, Tim J. P. Hubbard, Hugues Roest Crollius, Jane Rogers, and Derek L. Stemple. 2013. “The Zebrafish Reference Genome Sequence and Its Relationship to the Human Genome.” Nature.

Huang, Guo N., Jeffrey E. Thatcher, John McAnally, Yongli Kong, Xiaoxia Qi, Wei Tan, J. Michael DiMaio, James F. Amatruda, Robert D. Gerard, Joseph A. Hill, Rhonda Bassel-Duby, and Eric N. Olson. 2012. “C/EBP Transcription Factors Mediate Epicardial Activation during Heart Development and Injury.” Science.

Huang, Ying, Michael R. Harrison, Arthela Osorio, Jieun Kim, Aaron Baugh, Cunming Duan, Henry M. Sucov, and Ching-Ling Lien. 2013. “Igf Signaling Is Required for Cardiomyocyte Proliferation during Zebrafish Heart Development and Regeneration.” PloS One 8(6):e67266.

Ieda, Masaki, Takatoshi Tsuchihashi, Kathryn N. Ivey, Robert S. Ross, Ting Ting Hong, Robin M. Shaw, and Deepak Srivastava. 2009. “Cardiac Fibroblasts Regulate Myocardial Proliferation through B1 Integrin Signaling.” Developmental Cell 16(2):233–44.

Itou, J., I. Oishi, H. Kawakami, T. J. Glass, J. Richter, A. Johnson, T. C. Lund, and Y. Kawakami. 2012. “Migration of Cardiomyocytes Is Essential for Heart Regeneration in Zebrafish.” Development 139(22):4133–42.

Jopling, Chris, Eduard Sleep, Marina Raya, Mercè Martí, Angel Raya, and Juan Carlos Izpisúa Belmonte. 2010. “Zebrafish Heart Regeneration Occurs by Cardiomyocyte Dedifferentiation and Proliferation.” Nature 464(7288):606–9.

Kanisicak, Onur, Hadi Khalil, Malina J. Ivey, Jason Karch, Bryan D. Maliken, Robert N. Correll, Matthew J. Brody, Suh Chin J. Lin, Bruce J. Aronow, Michelle D. Tallquist, and Jeffery D. Molkentin. 2016. “Genetic Lineage Tracing Defines Myofibroblast Origin and Function in the Injured Heart.” Nature Communications 7.

Katz, Tamar C., Manvendra K. Singh, Karl Degenhardt, José Rivera-Feliciano, Randy L. Johnson, Jonathan A. Epstein, and Clifford J. Tabin. 2012. “Distinct Compartments of the Proepicardial Organ Give Rise to Coronary Vascular Endothelial Cells.” Developmental Cell.

Kikuchi, Kazu, Vikas Gupta, Jinhu Wang, Jennifer E. Holdway, Airon A. Wills, Yi Fang, and Kenneth D. Poss. 2011. “Tcf21+ Epicardial Cells Adopt Non-Myocardial Fates during Zebrafish Heart Development and Regeneration.” Development 138(14):2895–2902.

Kikuchi, Kazu, Jennifer E. Holdway, Andreas A. Werdich, Ryan M. Anderson, Yi Fang, Gregory F. Egnaczyk, Todd Evans, Calum A. Macrae, Didier Y. R. Stainier, and Kenneth D. Poss. 2010. “Primary Contribution to Zebrafish Heart Regeneration by Gata4(+) Cardiomyocytes.” Nature 464(7288):601–5.

Kim, Jieun, Qiong Wu, Yolanda Zhang, Katie M. Wiens, Ying Huang, Nicole Rubin, and Hiroyuki Shimada. 2010. “PDGF Signaling Is Required for Epicardial Function and Blood Vessel Formation in Regenerating Zebra FiSh Hearts.” 3–7.

Kinkel, Mary D., Stefani C. Eames, Louis H. Philipson, and Victoria E. Prince. 2010. “Intraperitoneal Injection into Adult Zebrafish.” Journal of Visualized Experiments : JoVE (42):3–6.

Lavine, Kory J., Kai Yu, Andrew C. White, Xiuqin Zhang, Craig Smith, Juha Partanen, and David M. Ornitz. 2005. “Endocardial and Epicardial Derived FGF Signals Regulate Myocardial Proliferation and Differentiation in Vivo.” Developmental Cell.

Lepilina, Alexandra, Ashley N. Coon, Kazu Kikuchi, Jennifer E. Holdway, Richard W. Roberts, C. Geoffrey Burns, and Kenneth D. Poss. 2006. “A Dynamic Epicardial Injury Response Supports Progenitor Cell Activity during Zebrafish Heart Regeneration.” Cell 127(3):607–19.

Li, Peng, Susana Cavallero, Ying Gu, Tim H. P. Chen, Jennifer Hughes, A. Bassim Hassan, Jens C. Brüning, Mohammad Pashmforoush, and Henry M. Sucov. 2011a. “IGF Signaling Directs Ventricular Cardiomyocyte Proliferation during Embryonic Heart Development.” Development.

Li, Peng, Susana Cavallero, Ying Gu, Tim H. P. Chen, Jennifer Hughes, A. Bassim Hassan, Jens C. Brüning, Mohammad Pashmforoush, and Henry M. Sucov. 2011b. “IGF Signaling Directs Ventricular Cardiomyocyte Proliferation during Embryonic Heart Development.” Development.

Limana, Federica, Chiara Bertolami, Antonella Mangoni, Anna Di Carlo, Daniele Avitabile, David Mocini, Pina Iannelli, Roberta De Mori, Carlo Marchetti, Ombretta Pozzoli, Carlo Gentili, Antonella Zacheo, Antonia Germani, and Maurizio C. Capogrossi. 2010. “Myocardial Infarction Induces Embryonic Reprogramming of Epicardial C-Kit+ Cells: Role of the Pericardial Fluid.” Journal of Molecular and Cellular Cardiology.

Männer, Jörg. 1999. “Does the Subepicardial Mesenchyme Contribute Myocardioblasts to the Myocardium of the Chick Embryo Heart? A Quail-Chick Chimera Study Tracing the Fate of the Epicardial Primordium.” Anatomical Record.

Mikawa, Takashi, and Robert G. Gourdie. 1996. “Pericardial Mesoderm Generates a Population of Coronary Smooth Muscle Cells Migrating into the Heart along with Ingrowth of the Epicardial Organ.” Developmental Biology.

Moerkamp, Asja T., Kirsten Lodder, Tessa Van Herwaarden, Esther Dronkers, Calinda K. E. Dingenouts, Fredrik C. Tengström, Thomas J. Van Brakel, Marie José Goumans, and Anke M. Smits. 2016. “Human Fetal and Adult Epicardial-Derived Cells: A Novel Model to Study Their Activation.” Stem Cell Research and Therapy.

Mosimann, Christian, Charles K. Kaufman, Pulin Li, Emily K. Pugach, Owen J. Tamplin, and Leonard I. Zon. 2011. “Ubiquitous Transgene Expression and Cre-Based Recombination Driven by the Ubiquitin Promoter in Zebrafish.” Development (Cambridge, England) 138(1):169–77.

Muraro, Mauro J., Gitanjali Dharmadhikari, Dominic Grün, Nathalie Groen, Tim Dielen, Erik Jansen, Leon van Gurp, Marten A. Engelse, Francoise Carlotti, Eelco J. P. de Koning, and Alexander van Oudenaarden. 2016. “A Single-Cell Transcriptome Atlas of the Human Pancreas.” Cell Systems 3(4):385-394.e3.

Porrello, Enzo R., Ahmed I. Mahmoud, Emma Simpson, Joseph A. Hill, James A. Richardson, Eric N. Olson, and Hesham A. Sadek. 2011. “Transient Regenerative Potential of the Neonatal Mouse Heart.” 331(February):1078–81.

Poss, Kenneth D., Lindsay G. Wilson, and Mark T. Keating. 2002. “Heart Regeneration in Zebrafish.” Science (New York, N.Y.) 298(5601):2188–90.

Rudat, Carsten, and Andreas Kispert. 2012. “Wt1 and Epicardial Fate Mapping.” Circulation Research.

Sánchez-iranzo, Héctor, María Galardi-castilla, Andrés Sanz-morejón, and Juan Manuel González-rosa. 2018. “Transient Fibrosis Resolves via Fibroblast Inactivation in the Regenerating Zebrafish Heart.”

Satoh, Akira, Gillian M. C. Cummings, Susan V. Bryant, and David M. Gardiner. 2010. “Neurotrophic Regulation of Fibroblast Dedifferentiation during Limb Skeletal Regeneration in the Axolotl (Ambystoma Mexicanum).” Developmental Biology.

Schnabel, Kristin, Chi Chung Wu, Thomas Kurth, and Gilbert Weidinger. 2011. “Regeneration of Cryoinjury Induced Necrotic Heart Lesions in Zebrafish Is Associated with Epicardial Activation and Cardiomyocyte Proliferation.” PLoS ONE.

Smart, Nicola, Sveva Bollini, Karina N. Dubé, Joaquim M. Vieira, Bin Zhou, Sean Davidson, Derek Yellon, Johannes Riegler, Anthony N. Price, Mark F. Lythgoe, William T. Pu, and Paul R. Riley. 2011. “De Novo Cardiomyocytes from within the Activated Adult Heart after Injury.” Nature.

Stelnicki, Eric J., Jeff Arbeit, Darrell L. Cass, Catherine Saner, Michael Harrison, and Corey Largman. 1998. “Modulation of the Human Homeobox Genes PRX-2 and HOXB13 in Scarless Fetal Wounds.” Journal of Investigative Dermatology.

Tessadori, Federico, Jan Hendrik van Weerd, Silja B. Burkhard, Arie O. Verkerk, Emma de Pater, Bastiaan J. Boukens, Aryan Vink, Vincent M. Christoffels, and Jeroen Bakkers. 2012. “Identification and Functional Characterization of Cardiac Pacemaker Cells in Zebrafish.” PLoS ONE 7(10).

Travers, Joshua G., Fadia A. Kamal, Jeffrey Robbins, Katherine E. Yutzey, and Burns C. Blaxall. 2016. “Cardiac Fibrosis: The Fibroblast Awakens.” Circulation Research 118(6):1021–40.

Vanlandewijck, Michael, Liqun He, Maarja Andaloussi Mäe, Johanna Andrae, Koji Ando, Francesca Del Gaudio, Khayrun Nahar, Thibaud Lebouvier, Bàrbara Laviña, Leonor Gouveia, Ying Sun, Elisabeth Raschperger, Markus Räsänen, Yvette Zarb, Naoki Mochizuki, Annika Keller, Urban Lendahl, and Christer Betsholtz. 2018. “A Molecular Atlas of Cell Types and Zonation in the Brain Vasculature.” Nature 554(7693):475–80.

Wang, Jinhu, Jingli Cao, Amy L. Dickson, and Kenneth D. Poss. 2015. “Epicardial Regeneration Is Guided by Cardiac Outflow Tract and Hedgehog Signaling.” Nature.

Weeke-Klimp, Alida, Noortje A. M. Bax, Anna Rita Bellu, Elizabeth M. Winter, Johannes Vrolijk, Josée Plantinga, Saskia Maas, Marja Brinker, Edris A. F. Mahtab, Adriana C. Gittenberger-de Groot, Marja J. A. van Luyn, Martin C. Harmsen, and Heleen Lie-Venema. 2010. “Epicardium-Derived Cells Enhance Proliferation, Cellular Maturation and Alignment of Cardiomyocytes.” Journal of Molecular and Cellular Cardiology.

Weinberger, Michael, Filipa C. Simões, Roger Patient, Tatjana Sauka-Spengler, and Paul R. Riley. 2020. “Functional Heterogeneity within the Developing Zebrafish Epicardium.” Developmental Cell 52(5):574-590.e6.

WHO. 2019. “Health Topics | Cardiovascular Diseases.” Cardiovascular Diseases.

van Wijk, Bram, Quinn D. Gunst, Antoon F. M. Moorman, and Maurice J. B. van den Hoff. 2012. “Cardiac Regeneration from Activated Epicardium.” PLoS ONE.

Wu, Chi Chung, Fabian Kruse, Mohankrishna Dalvoy Vasudevarao, Jan Philipp Junker, David C. Zebrowski, Kristin Fischer, Emily S. Noël, Dominic Grün, Eugene Berezikov, Felix B. Engel, Alexander van Oudenaarden, Gilbert Weidinger, and Jeroen Bakkers. 2015. “Spatially Resolved Genome-Wide Transcriptional Profiling Identifies BMP Signaling as Essential Regulator of Zebrafish Cardiomyocyte Regeneration.” Developmental Cell 36–49.

Yokoyama, Hitoshi, Tamae Maruoka, Akio Aruga, Takanori Amano, Shiro Ohgo, Toshihiko Shiroishi, and Koji Tamura. 2011. “Prx-1 Expression in Xenopus Laevis Scarless Skin-Wound Healing and Its Resemblance to Epimorphic Regeneration.” Journal of Investigative Dermatology.

Zangi, Lior, Kathy O. Lui, Alexander Von Gise, Qing Ma, Wataru Ebina, Leon M. Ptaszek, Daniela Später, Huansheng Xu, Mohammadsharif Tabebordbar, Rostic Gorbatov, Brena Sena, Matthias Nahrendorf, David M. Briscoe, Ronald A. Li, Amy J. Wagers, Derrick J. Rossi, William T. Pu, and Kenneth R. Chien. 2013. “Modified MRNA Directs the Fate of Heart Progenitor Cells and Induces Vascular Regeneration after Myocardial Infarction.” Nature Biotechnology.

Zhou, Bin, Leah B. Honor, Huamei He, Ma Qing, Jin Hee Oh, Catherine Butterfield, Ruei Zeng Lin, Juan M. Melero-Martin, Elena Dolmatova, Heather S. Duffy, Alexander Von Gise, Pingzhu Zhou, Yong Wu Hu, Gang Wang, Bing Zhang, Lianchun Wang, Jennifer L. Hall, Marsha A. Moses, Francis X. McGowan, and William T. Pu. 2011. “Adult Mouse Epicardium Modulates Myocardial Injury by Secreting Paracrine Factors.” Journal of Clinical Investigation.

